# CORONATINE INSENSITIVE 1-mediated repression of immunity-related genes in Arabidopsis roots is overcome upon infection with *Verticillium longisporum*

**DOI:** 10.1101/2024.03.27.586993

**Authors:** Louisa Ulrich, Johanna Schmitz, Corinna Thurow, Christiane Gatz

**Affiliations:** Department of Plant Molecular Biology and Physiology, Albrecht-von-Haller Institute for Plant Sciences, University of Göttingen, Julia-Lermontowa-Weg 3, 37077 Göttingen, Germany

**Keywords:** *Arabidopsis thaliana*, CORONATINE INSENSITIVE 1, jasmonoyl-isoleucine, root transcriptome, salicylic acid, split-root infections, SYSTEMIC ACQUIRED RESISTANCE DEFICIENT 1, *Verticillium longisporum*

## Abstract

*Verticillium longisporum* is a soil-borne fungal pathogen causing vascular disease predominantly in *Brassicaceae*. We have reported previously that the receptor of the plant defense hormone jasmonoyl-isoleucine (JA-Ile), CORONATINE INSENSITIVE 1 (COI1), is required in roots for efficient proliferation of the fungus in the shoot implicating a mobile root-borne signal that influences the outcome of the disease in shoots. To explore the underlying mechanism we compared the root transcriptome of *coi1* with the transcriptomes of three susceptible genotypes (wild-type, mutants deficient in JA-Ile and salicylic acid (SA) synthesis). At 10 days after infection, genes related to either xylem formation or plant immunity were induced independently of JA-Ile and SA. The biggest difference between the transcriptomes was due to 316 immunity-related genes that were pre-induced in *coi1*. Interfering with the expression of a subgroup of these genes partially suppressed the *coi1* phenotype. We therefore hypothesize that mobile defense compounds secreted into the xylem and being transported with the transpiration stream confer tolerance to the shoot. We furthermore report that 149 of the COI1-repressed genes are induced in WT upon infection reaching similar levels as in mock-treated *coi1*. The majority of these were not further induced in *coi1*, indicating that COI1 is required for infection-induced expression.

## Introduction

Plant roots are in close contact with a plethora of commensal, mutualistic and pathogenic microorganisms densely populating soil environments. The ascomycete fungus *Verticillium longisporum* is a soil-borne pathogen with a host range largely restricted to Brassicaceae. Mainly *Brassica napus* is an economically important host crop in Europe, to whose production *V. longisporum* poses an increasing threat (Depotter *et al*., 2016; Eynck *et al*., 2007). Moreover, *V. longisporum* is able to propagate in the model plant *Arabidopsis thaliana* (Iven *et al*., 2012; Ralhan *et al*., 2012; Reusche *et al*., 2012). *V. longisporum* penetrates roots and uses xylem vessels to spread systemically in its host. Infection with *V. longisporum* causes stunted growth, vein clearing, leaf chlorosis and premature senescence (Depotter *et al*., 2016; Ralhan *et al*., 2012).

Plant defense responses are activated upon recognition of microbe-associated molecular patterns (MAMPs) of potential pathogens by pattern recognition receptors, which are located in the plasma membrane (Boller and Felix, 2009). MAMPs are conserved molecules like e. g. flagellin, chitin or Necrosis and ethylene-inducing peptide 1 (Nep1)-like proteins (Chinchilla *et al*., 2006; Kaku *et al*., 2006; Oome *et al*., 2014). MAMP-induced signalling eventually activates the synthesis of plant defense hormones, i.e. salicylic acid (SA) or jasmonoyl-isoleucine (JA-Ile) in combination with ethylene. The corresponding hormone-mediated signalling pathways lead to massive transcriptional reprogramming to generate appropriate defense outputs (Wang *et al*., 2006; Zander *et al*., 2020).

After initial initiation of the SA and JA-Ile/ethylene pathways, only one of these dominates the defense response, depending on the type of pathogen invading (Spoel *et al*., 2007). Generalised, SA-mediated responses are deployed against biotrophic pathogens, which feed off living cells, while the JA-Ile/ethylene defense pathway is launched against necrotrophic pathogens, which kill the cells to feed on the remains (Glazebrook, 2005). *V. longisporum* can be regarded as a hemibiotroph as it starts off as a biotroph, spreading in the vascular system of the host plant without causing too much damage. Only later, before forming microsclerotia as resting structures, it starts to become necrotrophic and destroys the tissue (Depotter *et al*., 2016).

In petioles of *V. longisporum*-infected plants, SA and JA-Ile synthesis and the corresponding signalling pathways are activated, but the resulting defense mechanisms do not or only marginally influence fungal growth (Ralhan *et al*., 2012). Paradoxically, *V. longisporum*, as well as another soil-borne vascular ascomycete, *Fusarium oxysporum*, require the JA-Ile receptor CORONATINE INSENSITIVE 1 (COI1) for successful completion of their life cycles in Arabidopsis plants (Thatcher *et al*., 2009; Ralhan *et al*., 2012). In contrast to wild-type and the JA biosynthesis mutant *allene oxide synthase (aos)* plants, infected *coi1* mutants show less severe disease symptoms in shoots where lower fungal amounts are detected only at late stages of infection. Reciprocal grafts between *coi1* and WT revealed that COI1 is required in roots to cause susceptibility to *F. oxysporum* and *V. longisporum* in shoots (Thatcher *et al*., 2009; Ralhan *et al*., 2012). It has been suggested that COI1-mediated susceptibility against *F. oxysporum* is caused by fungal JA-Ile or a fungal JA-mimic that ‘hijacks’ COI1 to induce a mobile root-derived factor which promotes senescence in the shoot (Thatcher *et al*., 2009).

To explore the role of COI1 in *V. longisporum*-infected Arabidopsis plants, we performed an RNA-seq analysis of mock-treated and *V. longisporum*-infected roots of the susceptible genotypes WT, *aos*, and SA biosynthesis-impaired *sid2* (*salicylic acid induction-deficient 2*) and of the tolerant genotype *coi1*. Interestingly, *coi1* differed from the other genotypes by increased expression of immunity related genes already in mock-treated roots. Further analyses support the hypothesis that secreted defense compounds encoded by these pre-induced genes reach the shoot with the transpiration stream where they cause compromised propagation of the fungus despite wild-type like early colonization rates. Furthermore, we provide genetic evidence that lifting of the repressive function of COI1 after *V. longisporum* infection allows induction of immunity-related genes in roots.

## Materials and methods

### Plant and fungal material

All plants used in this article are in the *Arabidopsis thaliana* Col-0 background. Seeds from *aos* (SALK_017756) and *npr1-1* (N3726; Cao et al., 1997) were obtained from the Nottingham Stock Centre (NASC), Nottingham University, Nottingham, UK; *coi1* (SALK_035548) (Mosblech *et al*., 2011) was obtained from I. Heilmann (Martin-Luther-University, Halle, Germany); *sard1-1 cbp60g-1* (Zhang *et al*., 2010) was provided by Y. Zhang (UBC Vancouver, Canada) and *sid2-2* (Wildermuth *et al*., 2001) by F. M. Ausubel (Harvard University, Boston, USA). The c*oi1 sard1-1 cbp60g-1* triple mutant was generated through crossing of the respective above-mentioned genotypes. Primers for genotyping are listed in Supplementary Table S1. Infections were either done with *Verticillium longisporum* isolate Vl43 carrying an *enhanced GREEN FLOURESCENT PROTEIN* (*eGFP*) gene (Eynck *et al*., 2007) or with an untransformed Vl43 isolate provided by D. Cai (Christian-Albrechts-University, Kiel, Germany).

### Plant growth conditions and treatments

Cultivation and infection of plants were done as described (Ulrich *et al*., 2021). Plants were grown four 14 days on agar plates containing Murashige-Skoog-medium (MS) supplemented with 2% sucrose. After transfer onto a 1:1 mix of sand and twice-steamed soil on a thin layer of Seramis, cultivation was continued under short-day conditions at 120-140 μmol m^-2^ s^-1^. For genotyping, a single leaf was clipped from each plant during the first week of growth on the sand-soil mixture (Supplementary Fig. S1). After 14 days, plants were up-rooted. Roots were washed in tap water and allowed to rest in tap water as mock treatment or in a *V. longisporum* spore suspension for 45 minutes. Afterwards, plants were transferred into individual pots containing twice-steamed soil and kept for a final 10 to 21 days in short day conditions at 120-140 μmol photons m^-2^ s^-1^. Root and shoot material of one single plant were harvested for one biological replicate if not otherwise specified. For experiments shown in Fig. 4, seed batches of plants heterozygous for c*oi1* were initially placed on MS medium supplemented with 2% sucrose and 50 μM methyl jasmonate to identify plants homozygous for the *coi1* allele (Feys *et al*., 2001).

### Fungal culture and inoculation

Conidia from a glycerol stock solution were cultivated in liquid simulated xylem medium (SXM) (Hollensteiner *et al*., 2016) supplemented with 275 mg/L Cefotaxim for 7 days in a rotary shaker at 23°C and 90 rpm. Conidia were harvested by filtering through a fluted filter (Nucleo Bond folded filters, Macherey-Nagel, Dueren, Germany), washed in sterile tap water and their concentration was determined with a hemocytometer. Glycerol was added to a final concentration of 21.5%. The conidia infection stocks were initially stored at -20°C for 5 days and subsequently kept at -70°C until the day of inoculation. On inoculation day, conidia stocks were thawed, centrifuged for 8 min at 8000 rpm and resuspended in tap water to a final concentration of 5 x 10^5^ or 1 x 10^6^ spores/mL.

### Leaf area measurement

For disease phenotype analysis, photographs of individual plants were taken at 15 or 21 dpi. The surface area of the whole rosette was determined with the ‘BlattFlaeche’ Software (Datinf GmbH, Tuebingen, Germany) (Ralhan *et al*., 2012).

### Transcriptome analysis

Segregating populations resulting from *COI1 x coi1* and *AOS x aos* crosses were genotyped with primers specified in Table S1 (Ulrich *et al*., 2021). Twelve single homozygous roots of either WT*_aos_*, WT*_coi1_*, *aos, coi1* and *sid2* (mock-treated, *V. longisporum*-infected at 10 dpi) were combined for one replicate; replicates per genotype and treatment were obtained from four independent infection experiments. RNA extraction, mRNA sequencing and subsequent bioinformatic analysis was done as described (Ulrich *et al*., 2021) with one exception: reads aligned to the feature type “CDS” instead of “exon” were quantified. Enrichment of the SARD1 and CBP60g binding site ‘GAAATTT’ within the 1,000-bp upstream regions was performed as described previously (Berendzen *et al*., 2012; Zander *et al*., 2014) using the Cluster Analysis Real Randomization algorithm incorporated into Motif Mapper version 5.2.4.0.

### Quantitative real time (qRT)-PCR

RNA extraction, cDNA synthesis and qRT-PCR were performed as previously described (Ulrich *et al*., 2021). Calculations were done according to the 2^−Δ*C*T^ method (Livak and Schmittgen, 2001) using the *UBQ5* (AT3G62250) transcript as a reference. As shown in Fig. S2, *UBQ5* expression was independent from infections and genotypes. Primers used for qRT-PCRs are listed in Supplementary Table S2.

### Phytohormone measurements

Measurements of SA, SA glucoside, JA and JA-Ile were done as described (Yu *et al*., 2021).

### Statistical analysis

GraphPad Prism 10.0 (GraphPad Software, Inc., San Diego, CA) was used to conduct statistical analysis. Logarithmic (relative transcript levels) or linear (phytohormones, leaf area) values were subjected to either one- or two-way ANOVA followed by Tukey’s multiple comparison tests.

### Split-root experiment

Seedlings were grown vertically on Murashige-Skoog-medium (MS) supplemented with 2% sucrose. After 7 days, primary roots were cut below the first two lateral roots, and roots were allowed to grow for additional 14 days. Subsequently, each plant with two main roots of similar size was transferred onto a pot, in which two compartments were established with a small barrier made of overhead transparencies. After 14 days of cultivation on the 1:1 mix of sand and twice-steamed soil on a thin layer of Seramis (short-day conditions at a photon flux density of 120-140 μmol m^-2^ s^-1^), the two root systems were carefully placed into a 12 well microtiter plate containing either the spore suspension or tap water. Subsequently, plants were transferred to soil, with both root systems growing separately in two adjacent pots. After 12 days, roots of each pot were harvested separately (Supplementary Fig. S3).

### Transgenic plants and Western Blot analysis

To generate transgenic plants constitutively expressing SARD1, appropriate vectors were created via GATEWAY cloning (Invitrogen, Karlsruhe, Germany). The genomic *SARD1* region from the sequence encoding the translational start site to the codon for the last amino acid including introns was amplified using primers SARD1GWfwd and SARD1noStopGWrev (Table S3). The PCR product was inserted into the vector pDONR207 and subsequently recombined into pUBQ10GW3HAstrepII7 (Budimir *et al*., 2021) adding a sequence encoding a three times HA and StrepII C-terminal tag. The final construct pUBQ10-SARD1-3HAstrepII7 was used for *Agrobacterium tumefaciens*-mediated gene transfer into Col-0 wild-type and *sid2-2* mutant plants (Clough and Bent, 1998). To create empty vector (EV) control plants, Agrobacteria containing the plasmid pUBQ10GW3HAstrepII7 were used for plant transformation. Transgenic plants were characterised via BASTA (Bayer CropScience AG, Monheim, Germany) selection and Western Blot analysis was performed to assess SARD1-3xHA-StrepII protein levels in homozygous plants.

## Results

### Jasmonoyl-isoleucine and ISOCHORISMATE SYNTHASE 1 (ICS1)-derived salicylic acid do not influence colonization and transcriptional activity of V. longisporum-infected roots

In order to explore the JA-Ile-independent susceptibility-promoting function of COI1 in *V. longisporum*-infected roots, we subjected wild-type and *coi1* roots to transcriptome analysis after mock treatment and *V. longisporum* infection. In order to distinguish canonical JA-Ile/COI1-dependent genes from those that are dependent on COI1 but independent from AOS, *aos* was included in the experiment. Moreover, the *sid2* mutant, which is deficient in pathogen-induced SA synthesis, was analysed (Wildermuth *et al*., 2001). Since *coi1* and *aos* plants are male sterile (von Malek *et al*., 2002; Xie *et al*., 1998), we genotyped the seedlings obtained from heterozygous *AOS/aos* and *COI1/coi1* plants before infection experiments and used both respective outcrossed wild-type plants (WT*_aos_*, WT*_coi1_*) from each population as controls (Supplementary Fig. S1). RNA was derived from four experiments, each comprising combined shoots or roots from twelve plants per genotype and treatment. Harvest was at 10 days post infection (dpi) or 10 days after mock treatment. At this time point, comparable amounts of fungal DNA were detected in roots and shoots of all genotypes (Supplementary Fig. S4).

We obtained a first impression of the root transcriptomes (Supplementary Table S4) by principal component (PC) analysis (Fig. 1A). Samples of WT*_aos_*, WT*_coi1_*, *aos* and *sid2* were closely grouped. Differences on the PC1 axis occurred upon infection and affected all four genotypes in the same way. Hence, JA-Ile and SA do not strongly affect the transcriptional output of *V. longisporum*-infected roots.

**Fig. 1.**
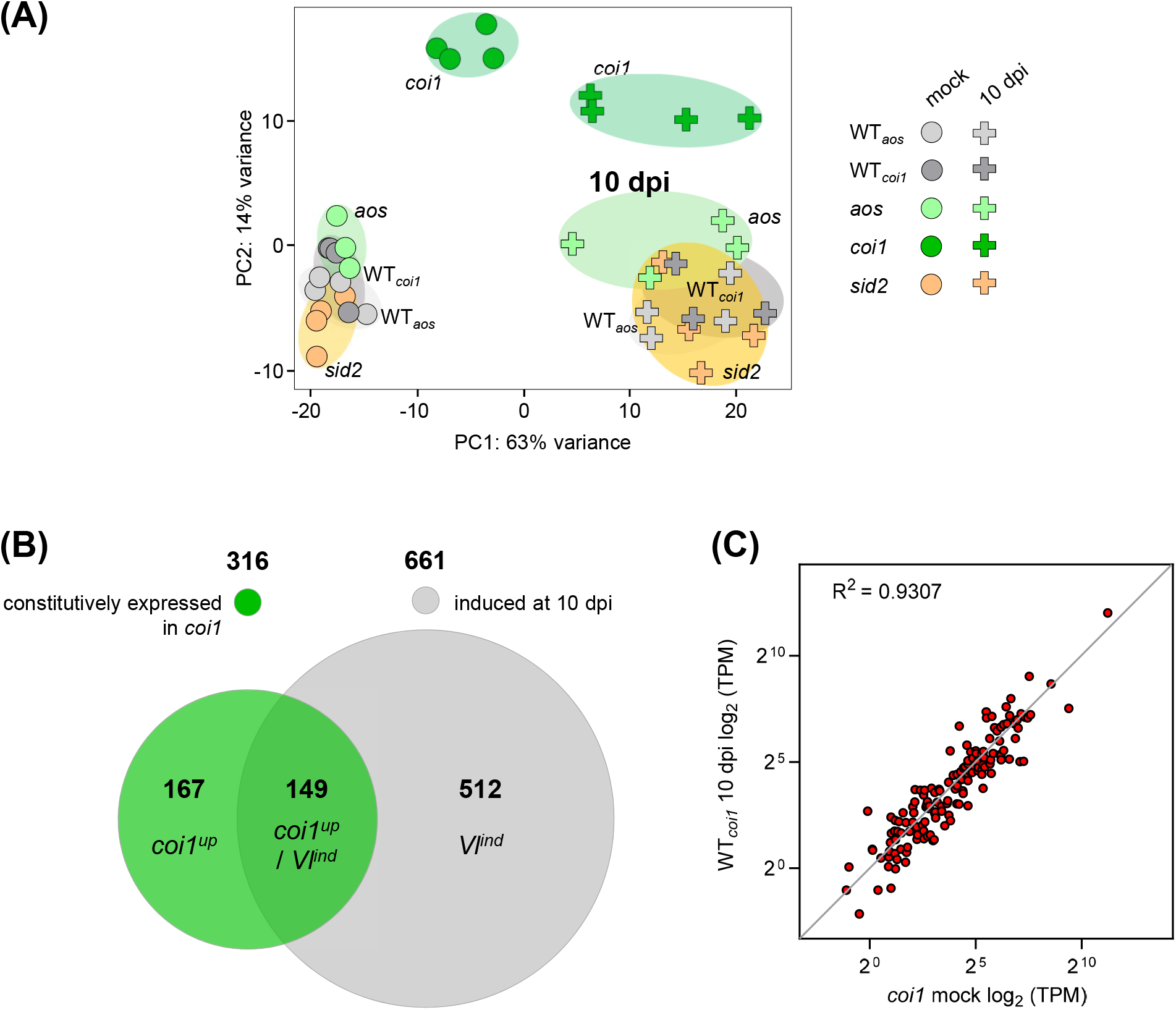
Analysis of normalized root transcriptome data of five genotypes after mock treatment and infection with *V. longisporum*. **(A)** Principal component analysis of the normalised root transcriptome data acquired by RNAseq analysis 10 days after mock treatment or inoculation with 1×10^6^ spores/mL eGFP-expressing *V. longisporum*. Biological replicates from four independent experiments are symbolised by circles (mock) or plus signs (10 dpi). For WT*_coi1_*, only three replicates were analysed for both mock and 10 dpi treatments. WT*_aos_* and WT*_coi1_*are the wild-types obtained from the segregating offspring of heterozygous *aos* and *coi1* plants. **(B)** Venn diagram showing the overlap between 316 genes constitutively upregulated in mock-treated *coi1* roots versus mock-treated WT*_aos_*, WT*_coi1_*, *aos* and *sid2* (> 2-fold, *p* < 0.05) and 661 genes induced in WT*_aos_*, WT*_coi1_*, *aos* and *sid2* at 10 dpi (> 2-fold, *p* < 0.05). Circles are drawn to scale with respect to the number of genes represented in each group. 167 genes constitutively expressed in *coi1* are referred to as ‘*coi1^up^*’, 149 genes constitutively expressed in *coi1* and induced upon *V. longisporum* infection are referred to as ‘*coi1^up^* / *Vl^ind^*’, 512 genes induced upon infection but not constitutively expressed in *coi1* are referred to as ‘*Vl^ind^*’. **(C)** Correlation analysis of the expression levels of 149 ‘*coi1^up^* / *Vl^ind^*’ genes in mock-treated *coi1* versus infected WT*_coi1_* plants. The blue line has a slope of 1. TPM, Transcripts Per Million.

After infection, 772 genes were induced in both WTs (> 2-fold, *p* < 0.05) (Supplementary Fig. S5A). In addition, 109 genes were induced only in WT*_aos_* and 176 genes were induced only in WT*_coi1_*. Most of these seemingly WT*_aos_*- or WT*_coi1_*-specific genes just closely missed the threshold. Gene Ontology (GO) term analysis showed that the 772 genes that were significantly induced in both WTs are associated with various processes in cell wall biogenesis and xylem development (Fig. 2, Supplementary Fig. S6). This is consistent with the notion that *V. longisporum* infections cause xylem hyperplasia, presumably to compensate for reduced water transport through the colonized vessels (Ralhan *et al*., 2012; Reusche *et al*., 2012). Both WTs show largely overlapping gene induction patterns with *sid2* after infection (Supplementary Figs S5B, S5C). Apparently, synthesis of SA through the isochorismate pathway (Wildermuth *et al*., 2001) does not play a major role for induction. Gene induction after infection in WT*_aos_* shows 72% overlap with gene induction in the *aos* mutant (Supplementary Fig. S5D), which is similar to the difference between WT*_aos_* and WT*_coi1_* (73%). This re-enforces the conclusion from the PCA analysis that had indicated a minor role of JA-Ile in gene expression at this time point.

**Fig. 2.**
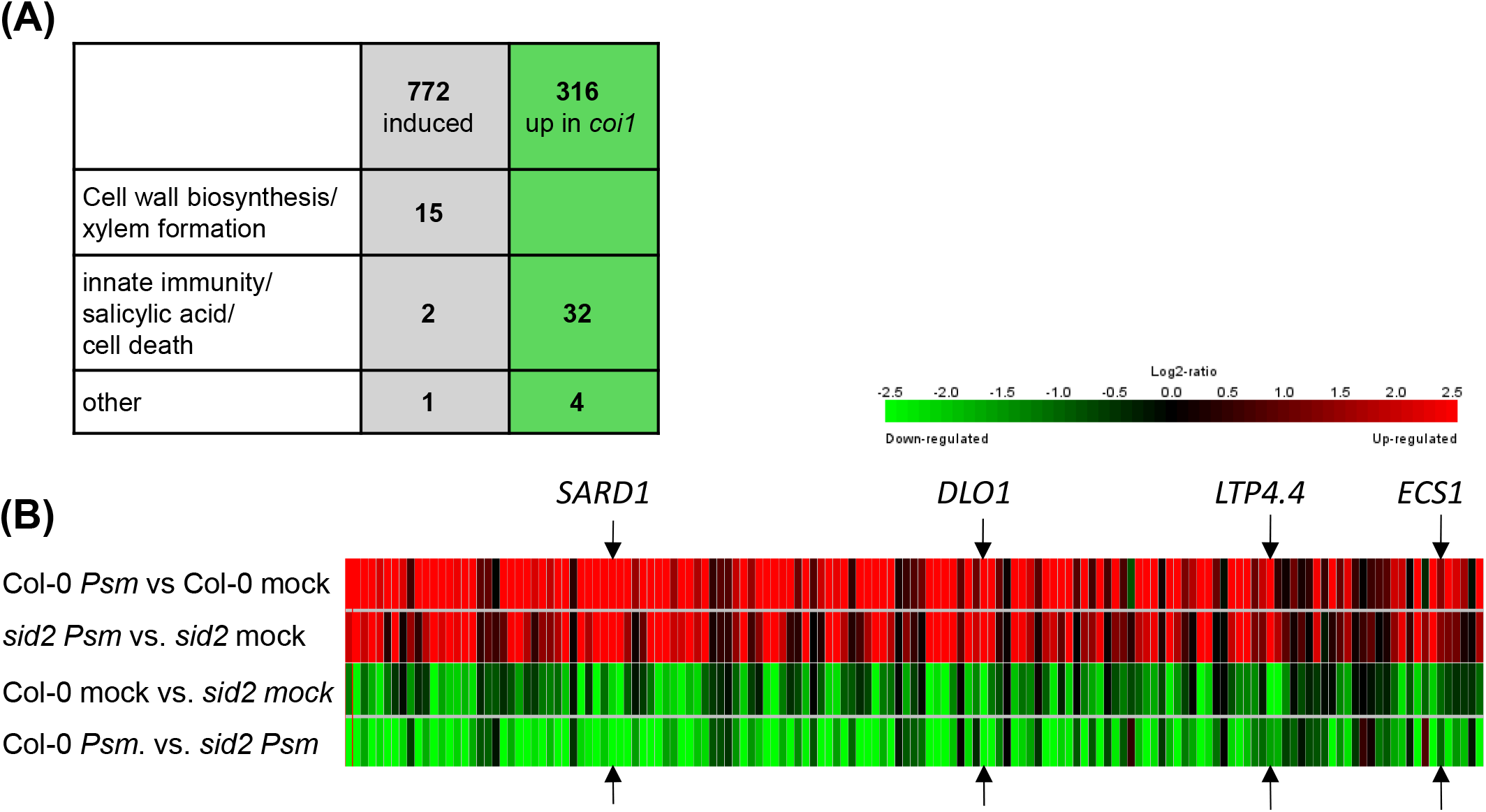
COI1 represses genes induced during systemic acquired resistance in the shoot. **(A)** Number of enriched GO terms in the group of 772 genes induced at 10 dpi in WT*_aos_* and WT*_coi1_* (grey) and in the group of 316 genes that were higher expressed in mock-treated *coi1* as compared to Wt, WT, *aos* and *sid2* (green). **(B)** 149 ‘*coi1^up^* / *Vl^ind^*’ genes are induced in systemic leaves of locally infected plants. *Arabidopsis thaliana* leaves were infected with *Pseudomonas syringae* pv. *maculicola* ES4326 (*Psm*) and distal leaves were harvested after 48 hours. Data are from https:\\genevestigator.com (At-00744(2), AT-00744(4), AT-00744(5), AT00744(6)). 149 ‘*coi1^up^* / *Vl^ind^*’ genes were arranged according to their fold induction in WT. The expression patterns of marker genes used in subsequent analysis are indicated by arrows.

Consistently, phytohormone measurements documented similar levels of SA and its glucoside SAG in infected and uninfected roots (Supplementary Fig. S7A). Likewise, JA levels stayed constant upon infection (Supplementary Fig. S7A). JA-Ile levels even decreased correlating with reduced transcript levels of *JA RESISTANT 1* (*JAR1*) (Supplementary Fig. S7B), which encodes an enzyme that catalyses the activation of JA to JA-Ile (Staswick and Tiryaki, 2004).

We continued our analysis with the most robustly differentially regulated genes, i.e. those that were induced upon infection (> 2-fold, *p* < 0.05) in all four susceptible genotypes, i.e. WT*_aos_*, WT*_coi1_*, *aos* and *sid2* (661 genes, Supplementary Table S5). 92 genes were down-regulated in these four genotypes after infection (> 2-fold, *p* < 0.05) (Supplementary Table S6).

### A subgroup of V. longisporum-induced genes is constitutively expressed in coi1

PC analysis revealed that only the transcriptome of *coi1* plants differed from those of the other genotypes. This difference became less after infection (Fig. 1A). The difference between mock-treated *coi1* roots as compared to the other mock-treated genotypes is mainly due to a set of 316 genes that was more highly expressed in *coi1* as compared to the other four genotypes (> 2-fold, *p* < 0.05) (Fig. 1B, Supplementary Table S7). Apart from *COI1*, only seven genes were lower expressed in mock-treated samples (Supplementary Table S8). The 316 COI1-repressed genes are associated with innate immune responses and SA-mediated programs like e.g. systemic acquired resistance (SAR) and programmed cell death (Fig. 2, Supplementary Fig. S8). The differences found after infection are due to 70 genes that were higher expressed (> 2-fold, p < 0.05) in *coi1* than in all other infected genotypes (Supplementary Table S9) and eleven genes with lower expression levels (Supplementary Table S10).

Interestingly, 149 of the 316 genes that were constitutively up-regulated in mock-treated *coi1* are also found in the 661 genes induced after infection in both WTs, *aos* and *sid2* (Fig. 1B, Supplementary Table S11). This group is called hereafter ‘*coi1^up^* / *Vl^ind^*’ for ‘constitutively up in *coi1*’ and ‘induced upon infection with *V. longisporum*’ in the other genotypes. Intriguingly, most of these 149 ‘*coi1^up^* / *Vl^ind^*’ genes were as highly expressed in mock-treated *coi1* plants as in infected WT*_coi1_* plants (Fig. 1C) with expression levels being not altered after infection. Analysis of the expression pattern of these genes in leaves (Zimmermann *et al*., 2004) revealed that many of these are induced in systemic leaves of plants infected with *Pseudomonas syringae* pv. *maculicola* ES4326 (*Psm*) (Fig. 2) (Bernsdorff *et al*., 2016). Similar to what we observed in roots, these genes are also induced in *sid2*. However, in contrast to the situation in roots, expression levels were much lower in *sid2* than in Col-0 indicating that SA serves as an aplifyer in shoots. The lack of enhanced SA levels in *V. longisporum*-infected roots (Supplementary Fig. S7A) is consistent with the missing amplification step.

The remaining 167 genes showing constitutive expression in *coi1* were not significantly (> 2-fold, *p* < 0.05) induced by *V. longisporum* in all other genotypes and are referred to as ‘*coi1^up^*’ (Fig. 1B, Supplementary Table S12). However, roughly one third of these was still significantly induced in at least one of the four genotypes.

The 512 genes, which were induced after infection in both WTs, *aos* and *sid2* and which were not significantly (> 2-fold, *p* < 0.05) higher expressed in mock-treated *coi1* as compared to the other mock-treated genotypes, are referred to as ‘*Vl^ind^*’ genes (Fig. 1B, Supplementary Table S13).

### Transcription factor SYSTEMIC ACQUIRED RESISTANCE DEFICIENT 1 (SARD1) is the master regulator of at least a subgroup of ‘coi1^up^ / Vl^ind^’ genes

Motif mapper analysis (Berendzen *et al*., 2012) showed that the motif ‘GAAATTT’ is significantly enriched in the promoters of the the 149 ‘*coi1^up^* / *Vl^ind^*’ genes but not in those of the 167 ‘*coi1^up^*’ genes (Fig. 3A). This motif is recognized by the partially redundant transcription factors SARD1 and CALMODULIN-BINDING PROTEIN 60-LIKE G (CBP60g) which are induced at 6 to 9 hours after MAMP perception in locally infected leaves and also later in systemic leaves (Bernsdorff *et al*., 2016; Wang *et al*., 2011; Zhang *et al*., 2010). They act as master regulators of SA and N-hydroxy-pipecolic acid biosynthesis (Sun *et al*., 2018; Wang *et al*., 2011; Zhang *et al*., 2010). Consistently, *SARD1* belongs to the 149 ‘*coi1^up^* / *Vl^ind^*’ genes. The expression patterns of *SARD1* and three further randomly chosen genes of the group of ‘*coi1^up^* / *Vl^ind^*’ genes (*LIPID TRANSFER PROTEIN4.4 (LTP4.4)* (AT5G55450), the cell wall-associated protein *ECOTYPE SPECIFIC 1 (ECS1)* (AT1G31580) (Aufsatz *et al*., 1998) and the SA catabolism gene *DMR6-LIKE OXYGENASE 1 (DLO1)* (AT4G10500)) are shown in Fig. 3B and Supplementary Fig. S9A. Transcript levels of these genes were as abundant in infected WTs, *aos* and *sid2* as in mock-treated *coi1* plants and were not further enhanced in *coi1* upon infection. Since their expression levels are amplified in shoots by SA (Fig. 2B), we tested whether induction might be supported by the SA receptor NON EXPRESSOR OF PATHOGENESIS RELATED 1 (NPR1), which was not the case (Fig. 3C, Supplementary Fig. S9B). We therefore conclude that transcription of these genes is not activated or amplified by basal SA levels or fungal SA mimics as found in leaves (Fig. 3C, Supplementary Fig. S9B).

**Fig. 3.**
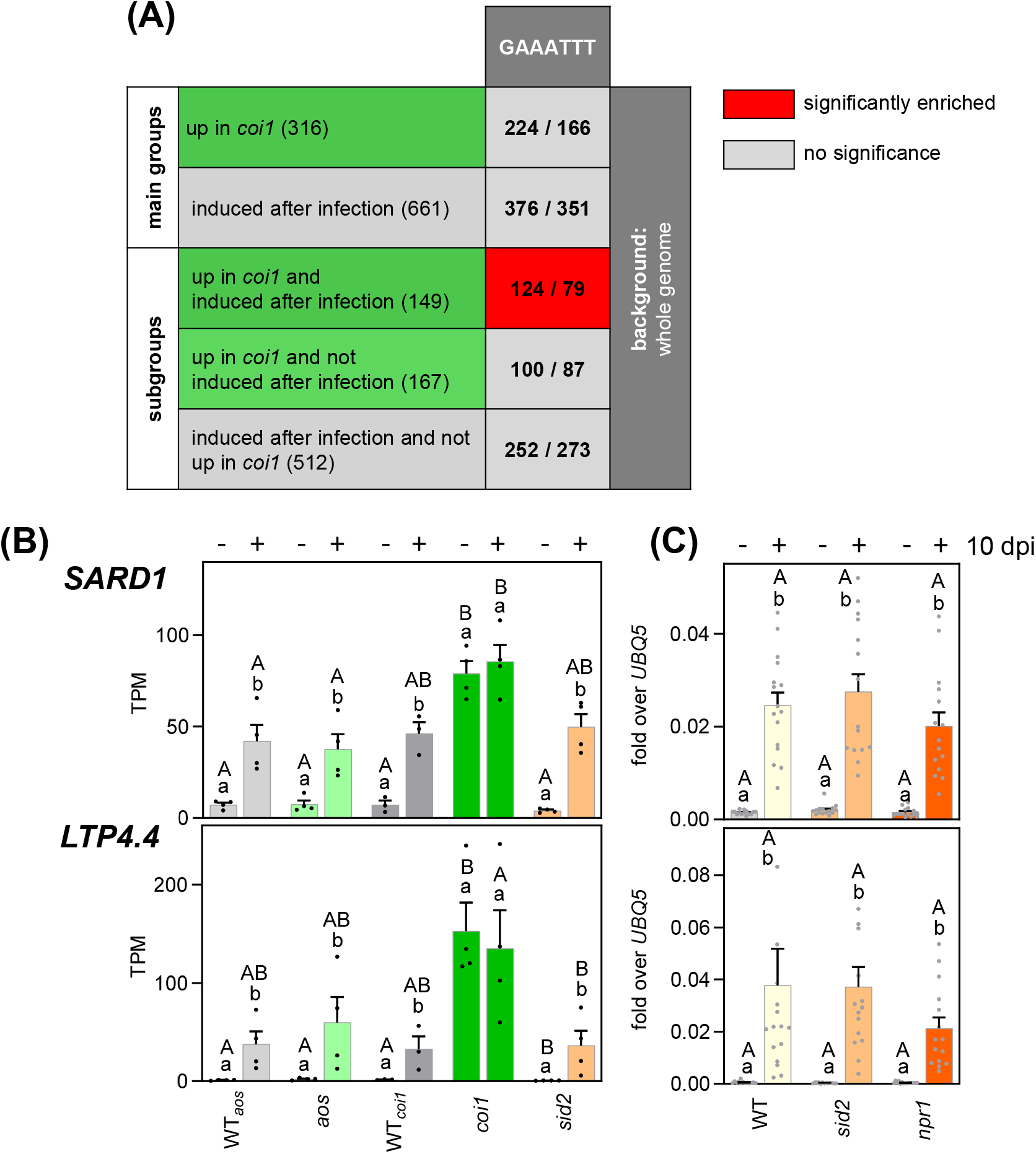
Identification of SARD1 as the potential master regulator of ‘*coi1^up^* / *Vl^ind^*’ genes. **(A)** Motif Mapper *cis* element analysis of the 316 genes de-repressed in *coi1* roots, the 661 genes induced in WT*_aos_*, WT*_coi1_*, *aos* and *sid2 at* 10 dpi, the 149 ‘*coi1^up^* / *Vl^ind^*’ genes, the 167 ‘*coi1^up^*’ genes, and the 512 ‘*Vl^ind^*’ genes according to Fig. 1B. **(B)** Relative *SARD1* and *LTP4.4* transcript levels as quantified by RNAseq analysis 10 days after mock treatment or inoculation with 1×10^6^ spores/mL eGFP-expressing *V. longisporum*. Bars are means of Transcripts Per Million (TPM) ± SEM of three to four biological replicates of each genotype, with each replicate representing twelve roots from one independent experiment. **(C)** Relative *SARD1* and *LTP4.4* transcript levels in WT, *sid2* and *npr1*, measured by qRT-PCR. RNA was extracted from roots at 10 days after mock treatment or infection with 1×10^6^ spores/mL *V. longisporum*. Bars are means ± SEM of thirteen to sixteen roots per genotype.For statistical analysis logarithmic values were subjected to a two-way ANOVA analysis followed by Tukey’s multiple comparison test; lowercase letters denote significant differences within each genotype between mock and 10 dpi (*p* < 0.05), uppercase letters denote significant differences between genotypes subjected to the same treatment (*p* < 0.05). WT*_aos_*and WT*_coi1_* are the two wild-type lines obtained from the segregating offspring of heterozygous *aos* and *coi1* seeds.

With SARD1 being a potential master regulator of the ‘*coi1^up^* / *Vl^ind^*’ genes, we asked the question whether pre-induction of *SARD1* in *coi1* might contribute to the tolerance phenotype of *coi1*. To this aim, we first determined if SARD1/CBP60g are required for gene induction in *coi1*. We chose to work with the *sard1 cbp60g* double mutant (Zhang *et al*., 2010) in order to avoid possible compensatory mechanisms of CBP60g in *sard1*. Indeed, expression of *LTP4.4*, *ECS1* and *DLO1* was reverted to WT levels in the *coi1 sard1 cbp60g* triple mutant (Fig. 4A). Likewise, the three marker genes were significantly less induced upon infection of *sard1 cbp60g* compared to WT at 10 dpi (Fig. 4A). Expression of exemplary genes belonging either to the ‘*coi1^up^*’ or the ‘*Vl^ind^*’ clusters was independent from SARD1/CBP60g (Supplementary Fig. S10).

**Fig. 4.**
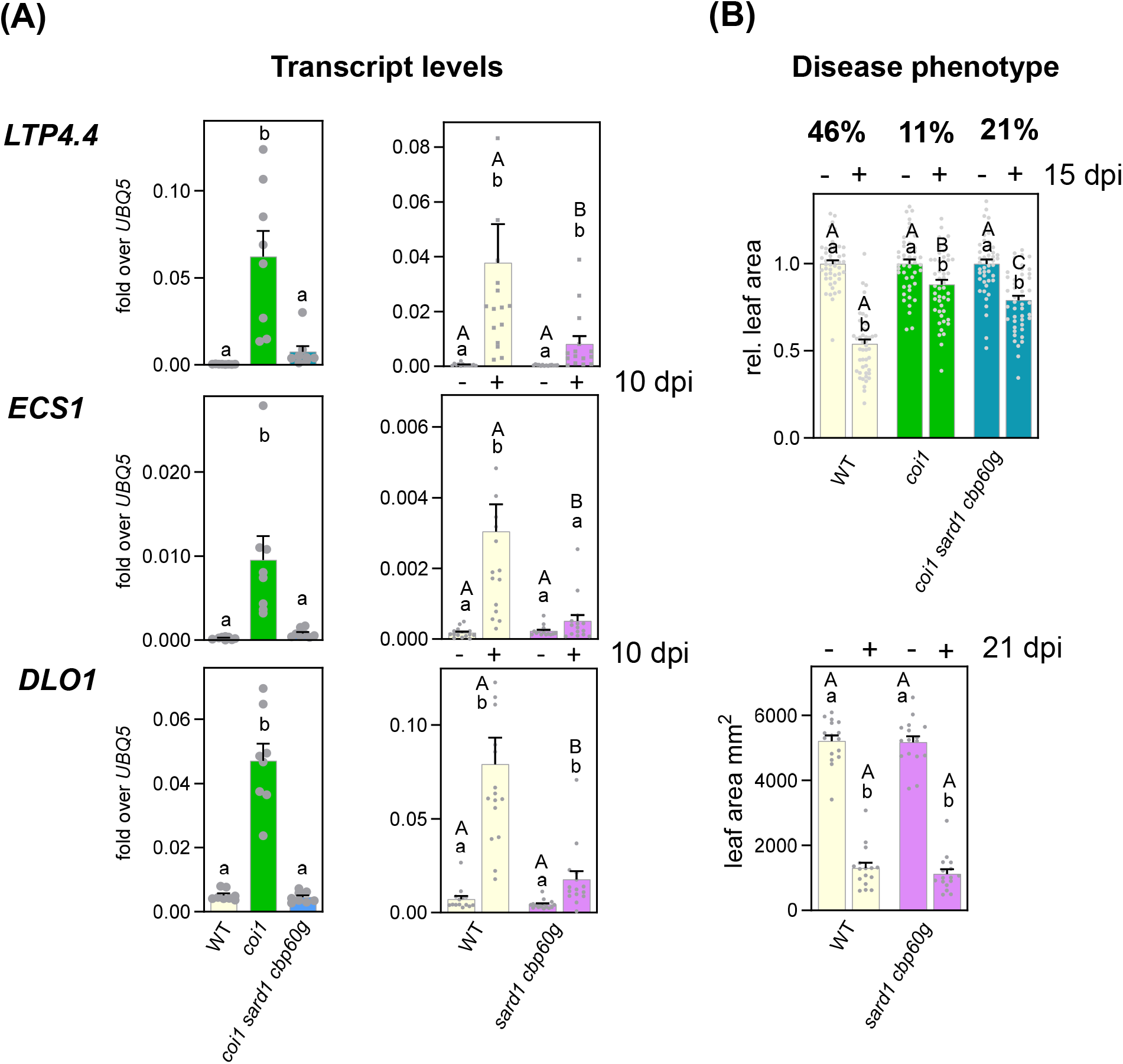
SARD1/CBP60g-regulated genes contribute to the tolerance phenotype of *coi1*. **(A)** Relative *LTP4.4, ECS1* and *DLO1* transcript levels, measured by qRT-PCR. RNA was extracted from mock-treated roots of the indicated genotypes (left panel; 8 plants per genotype) or after 10 days after mock treatment or infection with 1×10^6^ spores/mL *V. longisporum* of the indicated genotypes (16 plants per genotype and treatment). **(B)** Relative leaf area of WT, *coi1*, and *coi1 sard1 cbp60g* plants 15 days after mock treatment or infection with 5×10^5^ spores/mL eGFP-expressing *V. longisporum* (upper panel; 47-48 plants per treatment from three independent experiments with 15-16 plants per treatment each) or leaf area of WT and *sard1 cbp60g* 21 days after mock treatment or infection with 1×10^6^ spores/mL *V. longisporum.* Numbers above the upper panel indicate the per cent reduction of leaf area. For statistical analysis of the left panels in **(A)**, a one-way ANOVA was performed followed by Tukey’s multiple comparison test; letters denote significant differences between samples (*p* < 0.05). The remaining data were analysed by a two-way ANOVA followed by Tukey’s multiple comparison test; lowercase letters denote significant differences within each genotype between mock-treated and infected samples (*p* < 0.05), uppercase letters denote significant differences between genotypes subjected to the same treatment (*p* < 0.05). Logarithmic values were used in **(A).**

### Pre-induction of SARD1/CBP60g-dependent immunity-related genes might contribute to the tolerance phenotype observed in coi1

To address the question whether the constant activation of SARD1/CBP60g-dependent genes in *coi1* roots is the reason for the tolerance observed in *coi1* shoots, we infected *coi1 sard1 cbp60g* plants. While *coi1* plants showed 11% loss of leaf area at 15 dpi, WT plants showed 46% loss of leaf area and *coi1 sard1 cbp60g* plants were with 21% more similar to *coi1* than to WT plants (Fig. 4B). The intermediate phenotype might be due to those genes that are up-regulated in *coi1* independently of SARD1/CBP60g. Thus, these results suggest that pre-induction of the defense program in roots causes the tolerance observed in shoots. In contrast, we could not detect any significant difference between infected *sard1 cbp60g* and WT plants (Fig. 4B).

### SARD1 is not sufficient for the activation of ‘coi1^up^ / Vl^ind^’ genes in roots

Having identified SARD1/CBP60g as essential activators of at least a subfraction of ‘*coi1^up^* / *Vl^ind^*’ genes and to partially contribute to the *coi1* tolerance phenotype, we asked the question whether the increased *SARD1* expression that we observe in *coi1* renders plants more tolerant to *V. longisporum*. To this aim, we created transgenic lines that constitutively express *SARD1* (*SARD1 OX* lines). The construct contains the genomic *SARD1* sequence from the transcriptional start site and a C-terminal triple HA and a Strep-II tag under the control of the *UBIQUITIN10* (*UBQ10*) promoter. In order to distinguish between direct SARD1 effects and effects triggered by potential SARD1-induced SA synthesis, we introduced the *SARD1* construct not only into WT but also into *sid2* plants. For further analysis, we chose transgenic lines that had about twofold higher *SARD1* transcript levels compared to *V. longisporum*-infected plants (Supplementary Fig. S11A) which is in the same order of magnitude as *SARD1* transcript levels in *coi1* roots. Transcript and protein levels were similar in WT and *sid2* (Supplementary Fig. S11B).

As previously observed (Zhang *et al*., 2010), constitutive expression of *SARD1* resulted in moderate growth reduction due to the activation of the SA pathway (Supplementary Fig. S11C). In *SARD1 OX* lines, expression of target genes *LTP4.4* and *DLO1* was much more enhanced in shoots than in roots (Fig. 5). The influence of ectopically expressed SARD1 on *ECS1* was weak in both tissues. *ICS1* expression was twofold induced in shoots but not affected in roots. These data suggest that – in roots - COI1-mediated repression cannot be overcome by SARD1. In shoots, where this repression is not operational (Supplementary Fig. S12), elevated levels of SARD1 can be sufficient. Since enhanced expression of *SARD1* cannot mediate elevated expression of target genes in roots, increased tolerance is not expected.

**Fig. 5.**
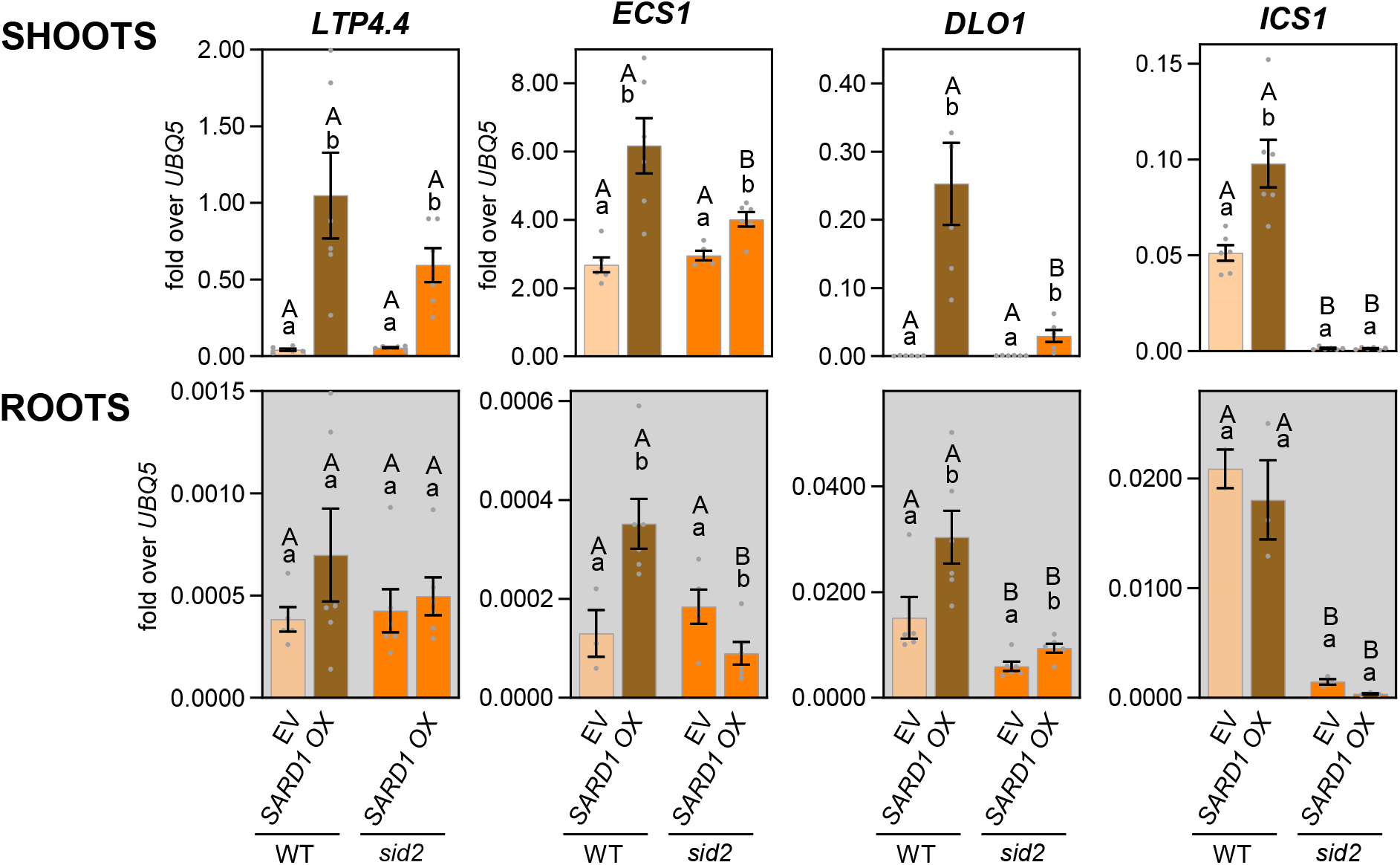
Overexpression of *SARD1* in roots does not lead to strong target gene activation in roots. Relative *LTP4.4, ESC1, DLO1* and *ICS1* transcript levels, measured by qRT-PCR. RNA was extracted from shoots or roots 10 days after mock treatment of *SARD1* overexpression lines (*SARD1ox*) and empty vector (EV) controls in WT and *sid2*. Bars are means ± SEM of three to six roots or shoots per line. For statistical analysis with logarithmic values, a two-way ANOVA was performed followed by Tukey’s multiple comparison test; lowercase letters denote significant differences within each genotype between mock and 10dpi (*p* < 0.05), uppercase letters denote significant differences between genotypes subjected to the same treatment (*p* < 0.05).

### Physically separated root systems communicate after V. longisporum infection

The observations that 149 ‘*coi1^up^* / *Vl^ind^*’ genes were expressed in infected WTs to approximately the same levels as in mock-treated *coi1* plants and that their expression in *coi1* was not altered upon infection (Fig. 3B) led to the hypothesis that they might be induced through a mechanism that involves inactivation of COI1. Assuming inactivation of COI1 as a possible induction mechanism, we reasoned that such a mechanism would require complete inactivation of COI1 upon infection, similar to what is observed in *coi1*. Since it is unlikely that the fungus infects every COI1 expressing cell, it has to be postulated that COI1 is inactivated systemically rather than only locally at the site of infection.

Systemic signalling in roots involves the synthesis of signal(s) travelling from the root to the shoot and back to the root (Ohkubo *et al*., 2017; Wang *et al*., 2019). In order to challenge the hypothesis of ‘systemic lifting of the repressive COI1 activity’, we performed experiments with split roots where the two halves of the root system were cultivated in different pots (Saiz-Fernandez *et al*., 2021) (Supplementary Fig. S3). Only one of the two main root systems was infected with *V. longisporum.* Potential fungal contamination of the mock-infected samples was excluded by qRT-PCR analysis of the fungal effector *HYQ44_006060* (GenBank: KAG7117937.1), which was first identified in *V. nonalfalfae* during colonization of hop plants (Flajsman *et al*., 2016) (Fig. 6B). Expression of *SARD1*, *LTP4.4* and *ECS1* was induced in the locally infected half, and – albeit to a lower extent – also in the mock-treated other half (Fig. 6C, Supplementary Figs S13, S14). Thus, the two halves of the root system seem to communicate. Transcription of *VASCULAR-RELATED NAC DOMAIN 7* (*VND7*), a transcription factor inducing the genetic program for xylem vessel formation (Yamaguchi *et al*., 2011), was only activated in the infected half of the root system (Fig. 6D). Thus, systemic signalling can lead to the induction of *SARD1* and its target genes.

**Fig. 6.**
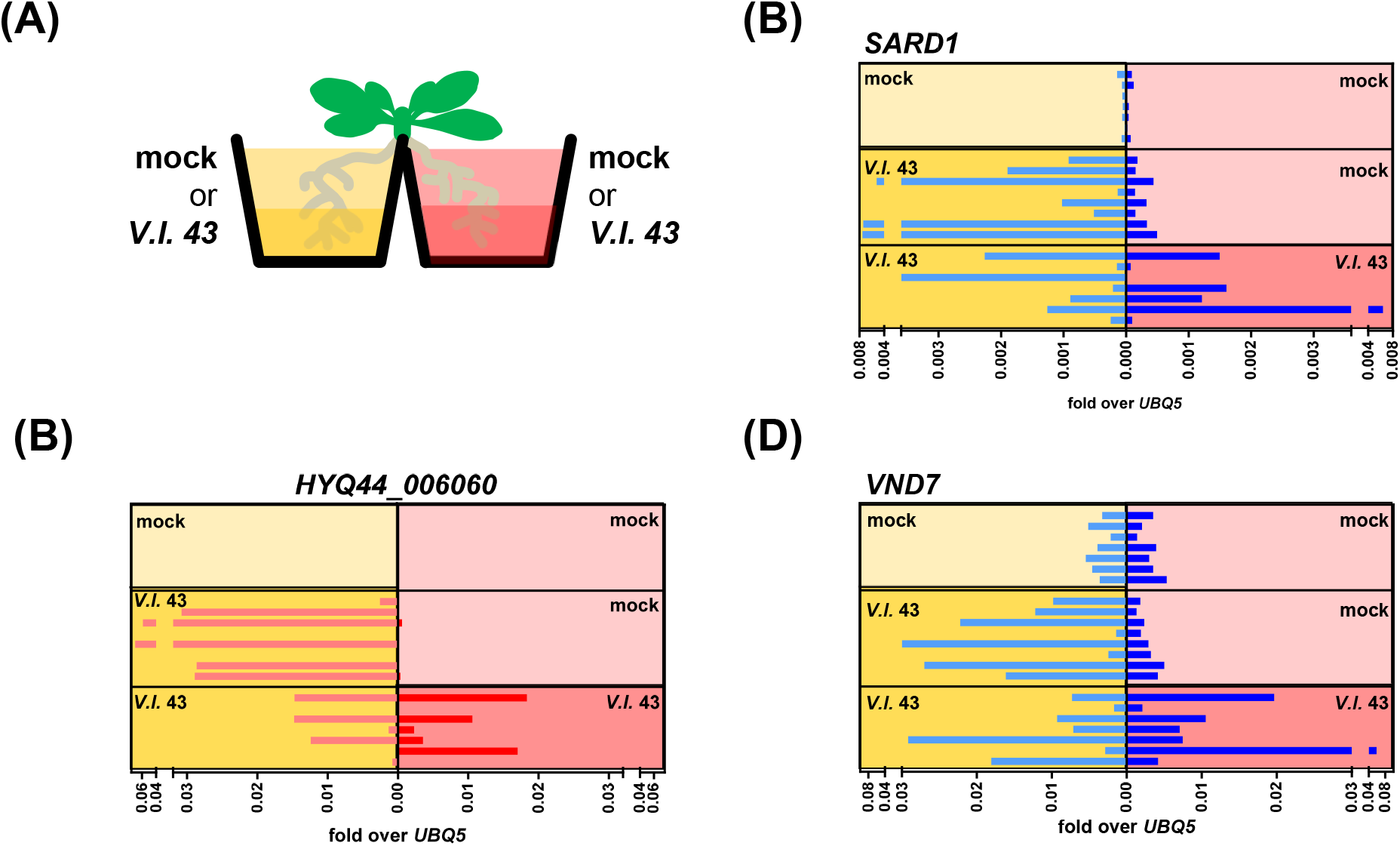
Expression analysis of *V. longisporum*-induced genes in physically separated roots. **(A)** Schematic outline of the split-root experiment. The two physically separated root systems were independently treated (mock/mock; *V. longisporum*/mock and *V. longisporum*/*V. longisporum*) RNA was extracted at 12 dpi. Relative expression of selected marker genes was determined by real time PCR, and values of the two root systems of each individual plant were plotted next to each other. **(B)** Relative transcript levels of the fungal effector *HYQ44_006060*. **(C)** Relative transcript levels of the ‘*coi1^up^* / *Vl^ind^*’ gene *SARD1*. **(D)** Relative transcript levels of the ‘*Vl^ind^*’ gene *VND7*.

## 3 Discussion

In this manuscript, we report the identification of genes that are differentially expressed in Arabidopsis roots upon *V. longisporum* infection at 10 dpi. Apart from genes associated with cell wall remodelling, a set of genes known to be upregulated upon MAMP perception in local and systemic leaves was induced independently of the major defense hormones SA or JA. In *coi1*, which exhibits a tolerance phenotype in shoots despite WT-like initial colonization of roots (Supplementary Fig. S4) (Ralhan *et al*., 2012), immunity-related transcript levels were already pre-induced to those levels that were found in WT upon induction. Elevated levels of extracellular root-to-shoot mobile defense proteins are thus candidates for conferring immunity. In *coi1*, transcripts were not further induced upon infection, suggesting a yet enigmatic role of COI1 for sensing the infection.

### In contrast to other root pathogens, V. longisporum does not lead to a hormonal imbalance in roots

Transcriptome analysis of *V. longisporum*-infected Arabidopsis roots has been performed before in two independent studies (Iven *et al*., 2012; Ulrich *et al*., 2021). Ulrich et al. (2021) did not observe any significant changes at the transcriptional level at 4 dpi when axenically grown plantlets were cultivated on agarose without any additional nutrients to avoid saprophytic growth of the fungus. In contrast, *Verticillium*-induced gene expression was detected when seedlings were continuously cultivated on synthetic medium with nutrients. Under these conditions, the fungus was able to colonize the cortex, but entry into the xylem seemed to be hampered (Iven *et al*., 2012; Reusche *et al*., 2014). At 3 dpi, expression of genes involved in tryptophan biosynthesis and tryptophan-derived secondary metabolism was induced leading to the synthesis of antimicrobial compounds (Iven *et al*., 2012). This was recently confirmed by the analysis of the translatome after infection (Froschel *et al*., 2021). Marker genes of the pathway for tryptophan-derived secondary metabolism like *CYP79b2* and *CYP79b3*, which were up-regulated in the axenic system (Iven *et al*., 2012), were not induced in soil-grown plants at 10 dpi (Supplementary Table S5). At this stage, fungal DNA was already detected in the shoot (Supplementary Fig. S4) indicating colonization of the xylem. GO term enrichment analysis of genes induced in soil-grown WT, *aos* and *sid2* unravelled preferential up-regulation of cell wall remodelling genes (Supplementary Fig. S6). Consistently, *VND7* coding for a master transcriptional regulator of *de novo* xylem formation (Reusche *et al*., 2012; Yamaguchi *et al*., 2011) was induced (Supplementary Table S5). In contrast, *VND7* was not induced when axenically grown plants were infected (Iven *et al*., 2012). The discrepancy of the transcriptional outputs under different conditions might be explained by the elicitation of specific defense responses upon colonization of the cortex, as observed early under axenic conditions at 3 dpi (Iven *et al*., 2012), while cell wall remodelling processes dominate the transcriptome upon establishment of the fungus in the xylem at 10 dpi.

The responses of *sid2* and *aos* were to the same degree different as the responses of the two WTs (Supplementary Fig. S5) indicating that SA and JA do not play a major role for the induction. This notion is supported by the fact that the abundance of transcripts of key biosynthesis enzymes of the pathways like *ICS1* (SA), *AOS* (JA) and *12-OXOPHYTODIEN REDUCTASE 3* (JA) were not or only slightly changed upon infection. *JAR1* expression was even significantly reduced correlating with reduced JA-Ile levels (Supplementary Fig. S7B).

Apart from the above mentioned transcriptomes of *V. longisporum*-infected roots, only a few transcriptomes of pathogen-infected Arabidopsis roots are published to our knowledge (*Fusarium oxysporum* (Lyons *et al*., 2015); *Phythophthora parasitica* (Attard *et al*., 2008); *Macrophomina phasaeolina* (Schroeder *et al*., 2019); *Plasmodiophora brassicae* (Irani *et al*., 2018)). Our transcriptome can be best compared to that of *Fusarium oxysporum*-infected roots, which was also from samples of soil-grown plants (Lyons *et al*., 2015). Tissue was collected at 6 dpi. Although *F. oxysporum* and *V. longisporum* are both vascular fungi, they seem to elicit quite distinct transcriptional responses. *F. oxysporum* elicits the activation of genes involved in JA synthesis and signalling. This might be either a consequence of root damage inflicted by *F. oxisporum*-derived phytotoxic compounds or, alternatively, of JA-Ile or JA-Ile mimics synthesized by the fungus initiating a feed forward loop leading to endogenous JA synthesis. Moreover, the activation of auxin-induced genes was reported, which might be a direct or indirect response to specific effector proteins. The strong cell wall remodelling response observed in *V. longisporum*-infected roots was not observed after *F. oxysporum* infection and is most likely initiated by a *V. longisporum*-specific effector.

Infection with the hemi-biotrophic pathogen *Phythophthora parasitica* induces SA- and JA-dependent signaling pathways during penetration (Attard *et al*., 2008). Later, during invasive growth, SA and JA pathways are de-activated and ethylene-mediated signaling predominates. Similar to what has been reported for the *V. longisporum* root transcriptome at 3 dpi (Iven *et al*., 2012), genes involved in tryptophan and camalexin synthesis are activated at all stages (Attard *et al*., 2014). *P. brassicae* induces the generation of abnormal tissue in the root, resulting in the formation of galls. Transcriptome analysis was performed with infected samples at 17, 20 and 24 dpi, when auxin-mediated processes are highly activated. The necrotrophic fungus *Macrophomina phasaeolina* triggers JA responses (Schroeder *et al*., 2019).

In conclusion, roots can respond with the activation of the SA- or JA pathway depending on the attacker. At 10 dpi with *V. longisporum*, transcriptome analysis of mutants deficient in SA or JA synthesis complemented by measurements of SA and JA levels showed that activation of SA and JA biosynthesis and signalling does not occur. In contrast, *V. longisporum* triggers both pathways in petioles at 15 dpi (Ralhan *et al*., 2012). Currently, it is not known, whether activation only in shoots is due to (i) a higher fungal load, (ii) different signals originating from the fungus when having reached the shoot, (iii) fungal effectors interfering with defense responses in the root or whether activation in only the shoot is due to (iv) a lower threshold for the elicitation of defense hormone biosynthesis as compared to roots.

### Induction of immunity-related genes might involve inactivation of the repressive function of COI1 at SARD1-dependent promoters

Despite the absence of increased SA or JA, a set of immunity-related genes was induced upon infection, including *SARD1* and its target genes. In locally infected leaves, expression of *SARD1* is triggered upon activation of MAMP-signalling (Sun *et al*., 2018), while in systemic leaves basal levels of SA, the SA receptor NPR1 and the hormone-like metabolite N-hydroxy-pipecolic acid are required for induction (Bernsdorff *et al*., 2016; Nair *et al*., 2021). In leaves, but not in roots, enhanced *SARD1* expression is sufficient to activate *ICS1* and the resulting enhanced SA levels further amplify gene expression in a feed forward loop (Fig. 5) (Sun *et al*., 2015).

It is noteworthy, that *SARD1* and its target genes were constitutively expressed in *coi1* and that expression was not significantly (> 2-fold, *p* < 0.05) further enhanced upon infection (Fig. 3, Supplementary Fig. S9). This has raised the hypothesis that – with regard to transcriptional activation – COI1 is important for sensing the infection. A possible scenario is that signals of either plant or fungal origin interfere with the repressive function of COI1, leading to the up-regulation of *SARD1*, which in turn can activate target genes in the absence of COI1 (Fig. 7). Since it is unlikely that each COI1-expressing cell is infected, inactivation of COI1 should occur systemically which would explain why constitutive expression levels in *coi1* are similar to expression levels observed after infection. Alternatively, MAMP signalling – although being active in roots only when co-inciding with damage (Zhou *et al*., 2020) - in infected cells might overcome COI1-mediated repression of SARD1 and high local SARD1 levels can subseqeuntly activate target genes by overriding the repressive COI1 effect. By coincidence, this high local expression would lead to similar expression levels as observed systemically in *coi1* when sampling the entire root system. However, in this scenario, we would expect further induction in locally infected cells of *coi1*, especially in view of pre-activation of signalling components similar to what is observed in SAR leaves (Supplementary Table S7). However, only 21 of the 149 ‘*coi1^up^* / *Vl^ind^*’ genes showed significantly enhanced transcript levels after infection in *coi1*. Hence, this model has to assume a dual role of COI1: first as a repressor, which affects all 316 genes, and second as a co-activator being required for high local expression of SARD1 which would be sufficient to activate at least a subgroup of the cluster of 149 ‘*coi1^up^* / *Vl^ind^*’ genes.

**Fig 7.**
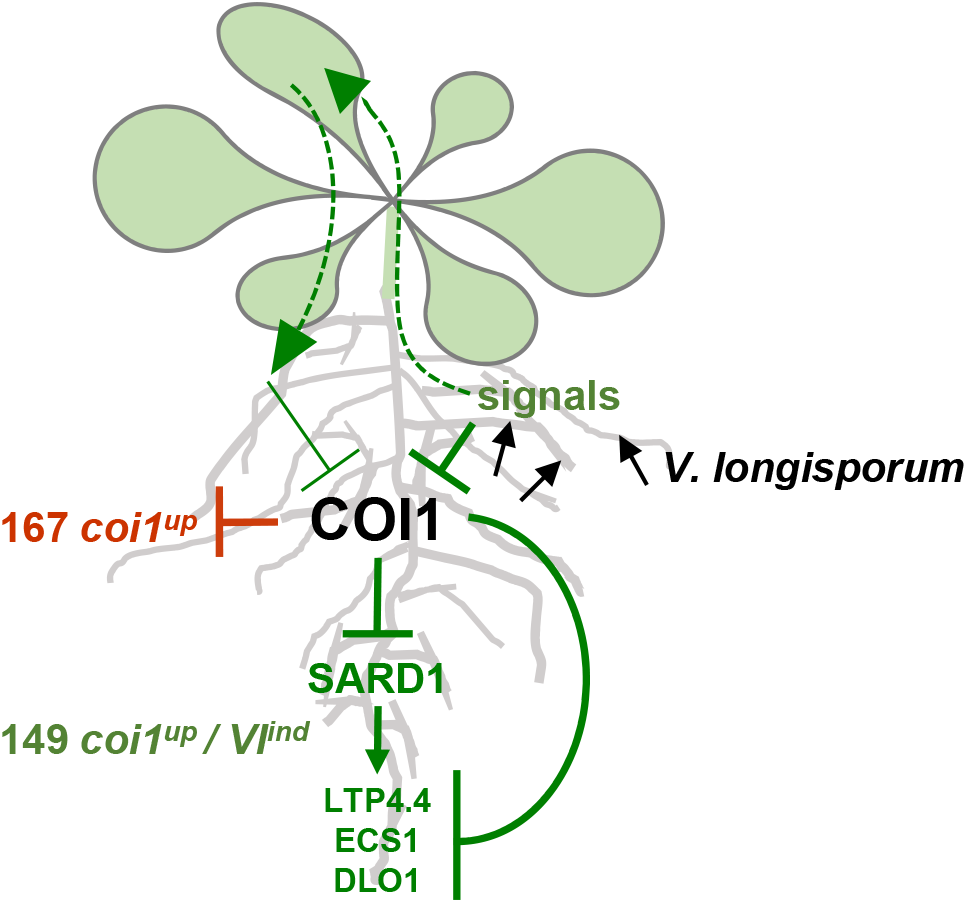
Working model explaining the activation of *V. longisporum*-inducible genes by inactivation of COI1. The working model suggests that *V. longisporum* infection generates a systemic signal that leads to inactivation of the repressive function of COI1 at primarily SARD1-regulated promoters (green), whereas SARD1-independent promoters (red) remain un-induced. The effect of the systemic signal is stronger near infection sites as after having travelled through the shoots.

The most direct way to differentiate between the two models would be to establish reporter lines under the control of promoters of genes like *SARD1*, *LTP4.4* and *ECS1*. However, microscopy of reporter gene expression and simultaneous localization of the GFP-tagged Verticillium in soil-grown roots is very likely challenging. Therefore, we decided to perform a split-root experiment which allows to test for systemic induction of several marker. Indeed, we found weak induction of *SARD1* and its target genes *LTP4.4* and *ECS1* in systemic roots. Since increased SARD1 levels as in *SARD1 OX* lines cannot activate target gene expression, while similarily increased SARD1 levels in *coi1* can do so, we assumed that COI1 needs to be inactivated to allow SARD1 function at target promoters. In the split-root experiment, weak induction of *SARD1* led to increased expression of target genes. It is therefore tentatively concluded that systemic signalling through the root-shoot-root path can partially overcome the repressive function of COI1 at SARD1-regulated promoters (Fig. 7). Chromatin immunoprecipitation experiments with leaves have identified SARD1 at its own promoter and at 80 of the 149 target genes (Sun *et al*., 2015).This effect might be much more pronounced in the vicinity of infection sites. Further research is required for understanding the role of COI1 at SARD1-regulated promoters.

### Genes constitutively de-repressed in coi1 might cause the tolerance phenotype of the shoot

The original aim of the study was to obtain information on the mechanism of COI1-mediated tolerance against *V. longisporum*. Grafting studies had unravelled that COI1 but not AOS has to be expressed in roots for successful completion of the life cycles of the two vascular pathogens *V. longisporum* and *F. oxysporum* in shoots (Ralhan *et al*., 2012; Thatcher *et al*., 2009). The tolerance phenotype is not due to up-regulation of the SA pathway as revealed by the analysis of *coi1* plants with reduced SA levels as in *coi1 nahG* or *coi1 sid2* (Thatcher *et al*., 2009; Ulrich *et al*., 2021). Since initial colonization of the shoot is similar in *coi1* as in WT and *aos* (Supplementary Fig. S4) (Ralhan *et al*., 2012; Thatcher *et al*., 2009) local defense responses do not explain the phenomenon. It rather has to be postulated that mobile signals mediating either susceptibility or resistance are misregulated in *coi1*. Since several *F. oxysporum* strains are able to produce JA-Ile (Oliw and Hamberg, 2019), it has been hypothesized that fungal JA-Ile or JA-Ile mimics “hijack” COI1 to initiate processes in the root that promote senescence in the shoot (Thatcher *et al*., 2009). In senescent tissue, the fungus might profit from weakened cell walls and transport of metabolites from mesophyll cells to the vascular system. This scenario is unlikely for *V. longisporum*, because potential fungal JA-Ile or JA-Ile mimics should induce JA marker genes like e.g. *JAZs*, *THIONIN2.1* (AT1G72260) and *PLANT DEFENSIN 1.2* (AT5G44420) in *aos* roots, which is not the case (Supplementary Table S5).

Only sixteen genes were at most threefold lower expressed in *coi1* than in *aos* and the other susceptible genotypes (Supplementary Tables S8, S10). None of them gives rise to an obvious hypothesis on their potential involvement in susceptibility. Still, the hypothetical susceptibility-promoting signal might be generated earlier or later than at 10 dpi and/or might not be detected at the transcript level.

Interestingly, 316 genes related to innate immunity, SAR and cell death are constitutively up-regulated in c*oi1* roots (Fig. 2, Supplementary Fig. S8), most of them coding for signalling proteins like receptor kinases and transcription factors. Still, highly expressed defense compounds like e.g. extracellular chitinases (AT4G01700), hydrolases (AT4G19720), peroxidases (AT2G38390) or other antimicrobial compounds like germin-like proteins (AT5G39110, AT5G39120, AT5G39130, AT5G39150, AT5G39160, AT5G39180, AT5G39190) (Pei *et al*., 2019; Pei *et al*., 2020) might be secreted into the xylem vessels so that they accumulate in the shoot when being permanently produced in the root. In *coi1 sard1 cbp60g*, some of these are likely to be less expressed which explains the partially reduced tolerance of this mutant (Fig. 4B). The *sard1 cbp60g* double mutant was not more susceptible to the fungus as compared to WT (Fig. 4B) enforcing the idea that pre-induction is important. This is different from what has been reported for infections with *V. dahliae*, which were slightly but significantly more successful on *sard1 cbp60g* (Qin *et al*., 2018). Our data set can be regarded as support of the notion that pre-induction of immunity related genes in *coi1* leads to the accumulation of root-to-shoot mobile defense compounds that counteract fungal growth in the shoot.

## Abbreviations

AOS: ALLENE OXIDE SYNTHASE
CBP60g: CALMODULIN-BINDING PROTEIN 60-LIKE G
COI1: CORONATINE INSENSITIVE 1; ‘*coi1^up^*’: cluster of genes up-regulated in *coi1*; ‘*coi1^up^* / *Vl^ind^*’: cluster of genes up-regulated in *coi1* and induced upon infection with *V. longisporum*
DLO1: DMR6-LIKE OXYGENASE 1
ECS1: ECOTYPE SPECIFIC 1
eGFP: enhanced GREEN FLOURESCENT PROTEIN
ERF54: ETHYLENE RESPONSE FACTOR 54
GO: Gene Ontology
ICS1: ISOCHORISMATE SYNTHASE 1
JA: jasmonic acid
JA-Ile: jasmonoyl-isoleucine
JAR1: JASMONATE RESISTANT 1
JAZ: JASMONATE ZIM-domain
LTP4.4: LIPID TRANSFER PROTEIN 4.4
MAMP: microbe-associated molecular pattern
NPR1: NON EXPRESSOR OF PATHOGENESIS RELATED 1
PC: principal component
SA: salicylic acid
SID2: SALICYLIC ACID INDUCTION DEFICIENT 2, SAR: systemic acquired resistance
SARD1: SYSTEMIC ACQUIRED RESISTANCE DEFICIENT 1
UBQ: UBIQUITIN; ‘*Vl^ind^*’: cluster of genes induced upon infection with *V. longisporum*, not up-regulated in *coi1*
VND7: VASCULAR-RELATED NAC DOMAIN 7
WT*_aos_*: segregating wild-type plants from a cross between wild-type and *aos*
WT*_coi1_*: segregating wild-type plants from a cross between wild-type and *coi1*

## Supplementary data

The following supplementary data are available at JXB online.

**Supplementary Table S1.**
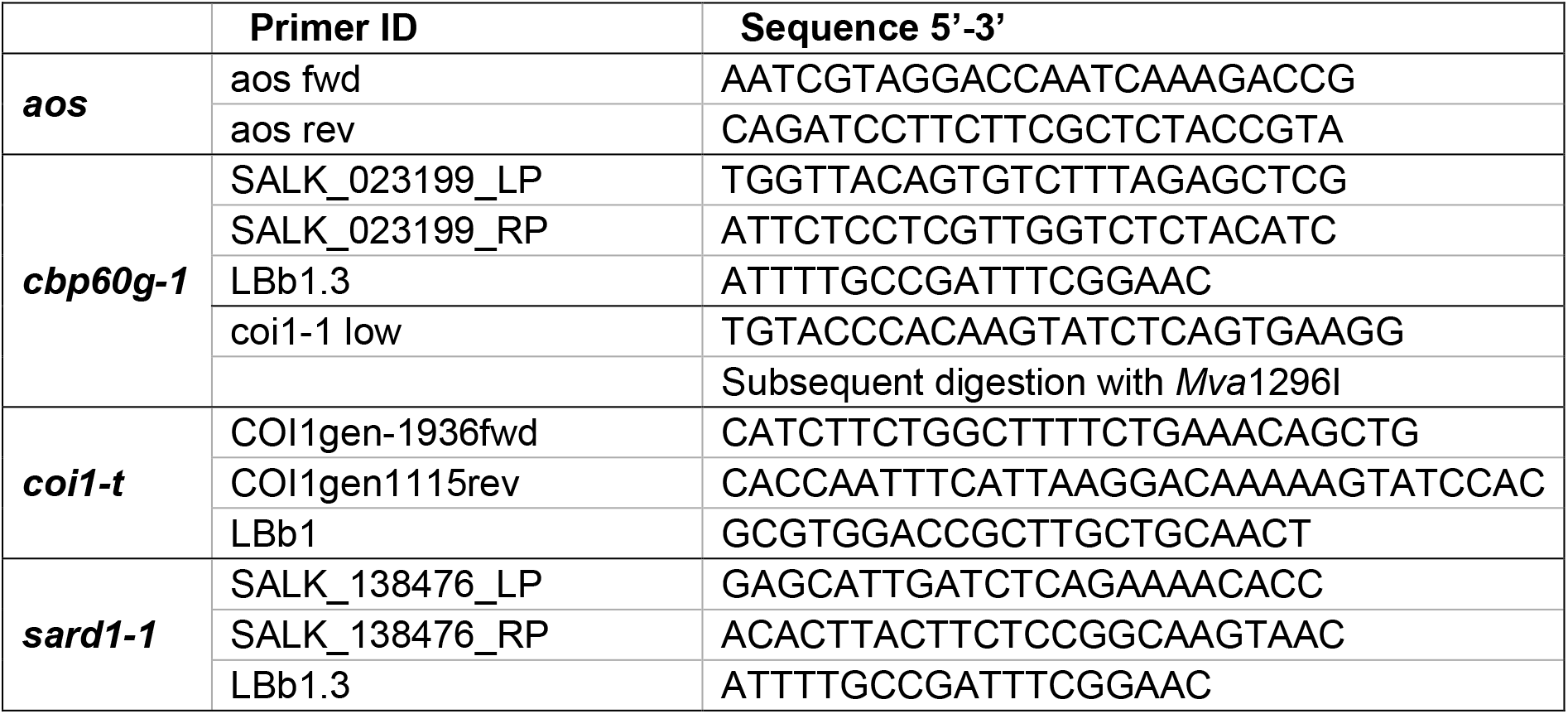
Primers for genotyping.

**Supplementary Table S2.**
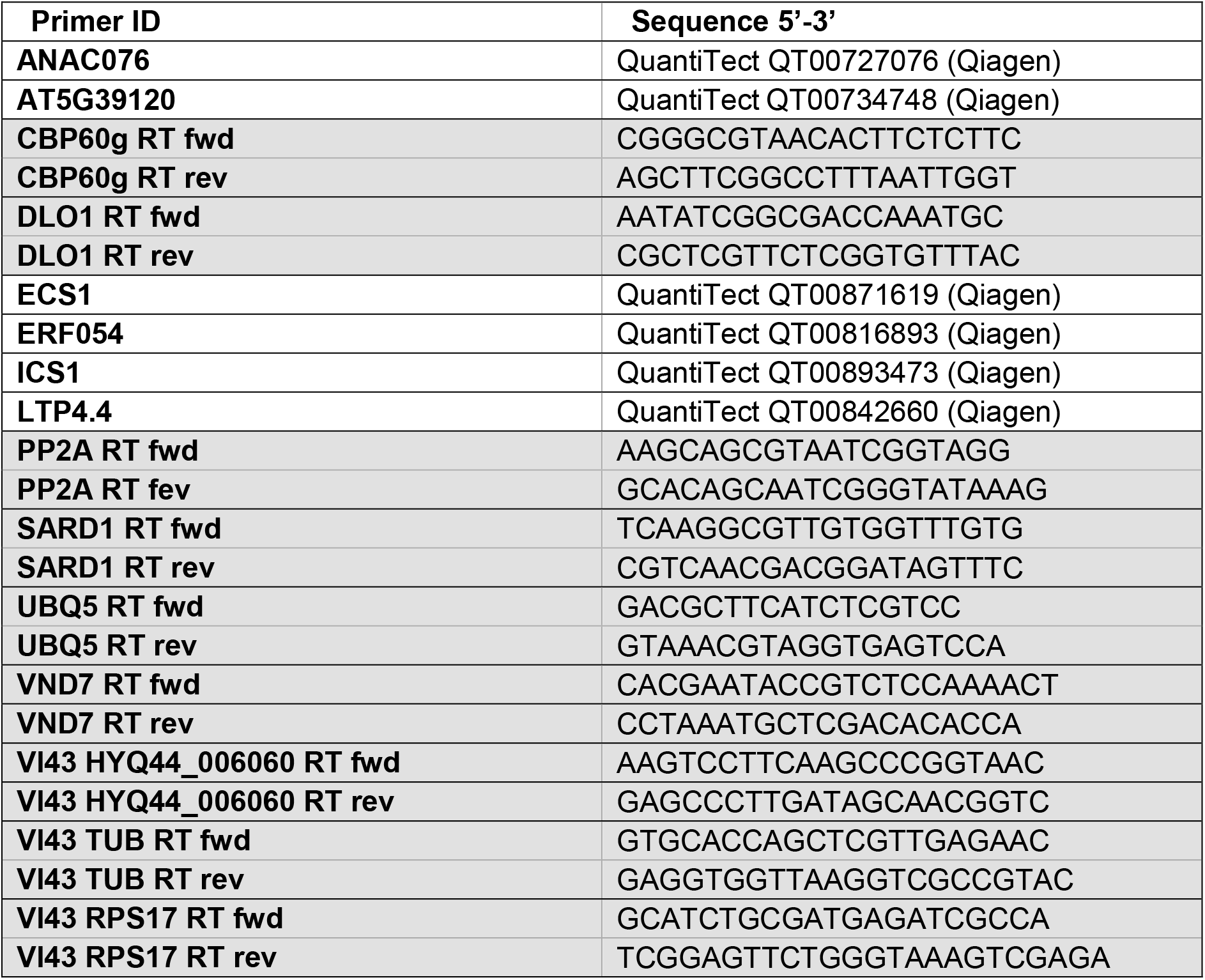
Primers for qRT-PCR.

**Supplementary Table S3.**
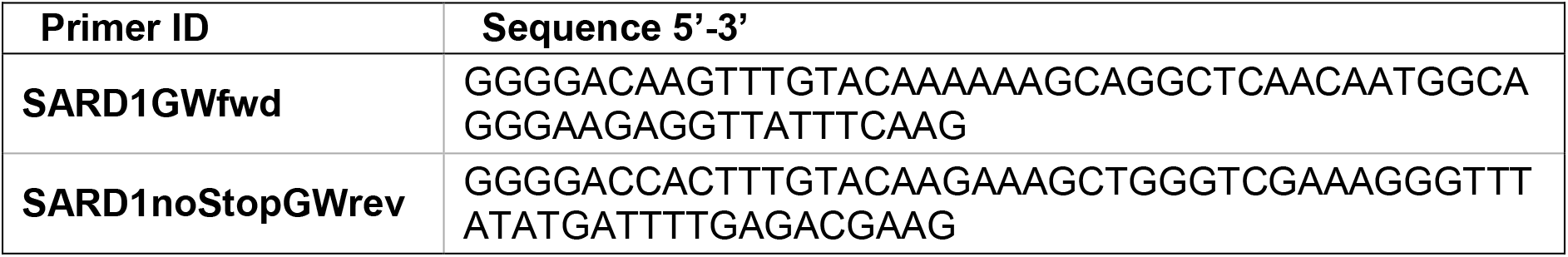
Primers for Cloning.

*Table S4*. Transcriptome data of mock-treated and *V. longisporum*-infected Arabidopsis plants (WT*_aos_*, WT*_coi1_*, *aos, coi1* and *sid2*).

*Table S5*. Expression data of 661 genes induced upon *V. longisporum* infection in WT*_aos_*, WT*_coi1_*, *aos* and *sid2*.

*Table S6*. Expression data of 92 genes down-regulated upon *V. longisporum* infection in WT*_aos_*, WT*_coi1_*, *aos* and *sid2*.

*Table S7*. Expression data of 316 genes with elevated expression levels in mock-treated *coi1* as compared to mock-treated WT*_aos_*, WT*_coi1_*, *aos* and *sid2*.

*Table S8*. Expression data of seven genes with lower expression levels in mock-treated *coi1* as compared to mock-treated WT*_aos_*, WT*_coi1_*, *aos* and *sid2*.

*Table S9:* Expression data of70 genes with elevated expression levels in infected *coi1* as compared to infected WT*_aos_*, WT*_coi1_*, *aos* and *sid2*.

*Table S10:* Expression data of twelve genes with lower expression levels in infected *coi1* as compared to infected WT*_aos_*, WT*_coi1_*, *aos* and *sid2*.

*Table S11*. Expression data of 149 ‘*coi1^up^* / *Vl^ind^*’ genes (induced upon *V. longisporum* infection in WT*_aos_*, WT*_coi1_*, *aos* and *sid2*, and up-regulated in mock-treated *coi1*).

*Table S12*. Expression data of 167 ‘*coi1^up^*’ genes (up-regulated in mock-treated *coi1*, but not induced upon *V. longisporum* infection).

*Table S13*. Expression data of 512 ‘*Vl^ind^*’ genes (induced upon *V. longisporum* infection in WT*_aos_*, WT*_coi1_*, *aos* and *sid2*, but not increased in mock-treated *coi1*).

**Fig. S1.**
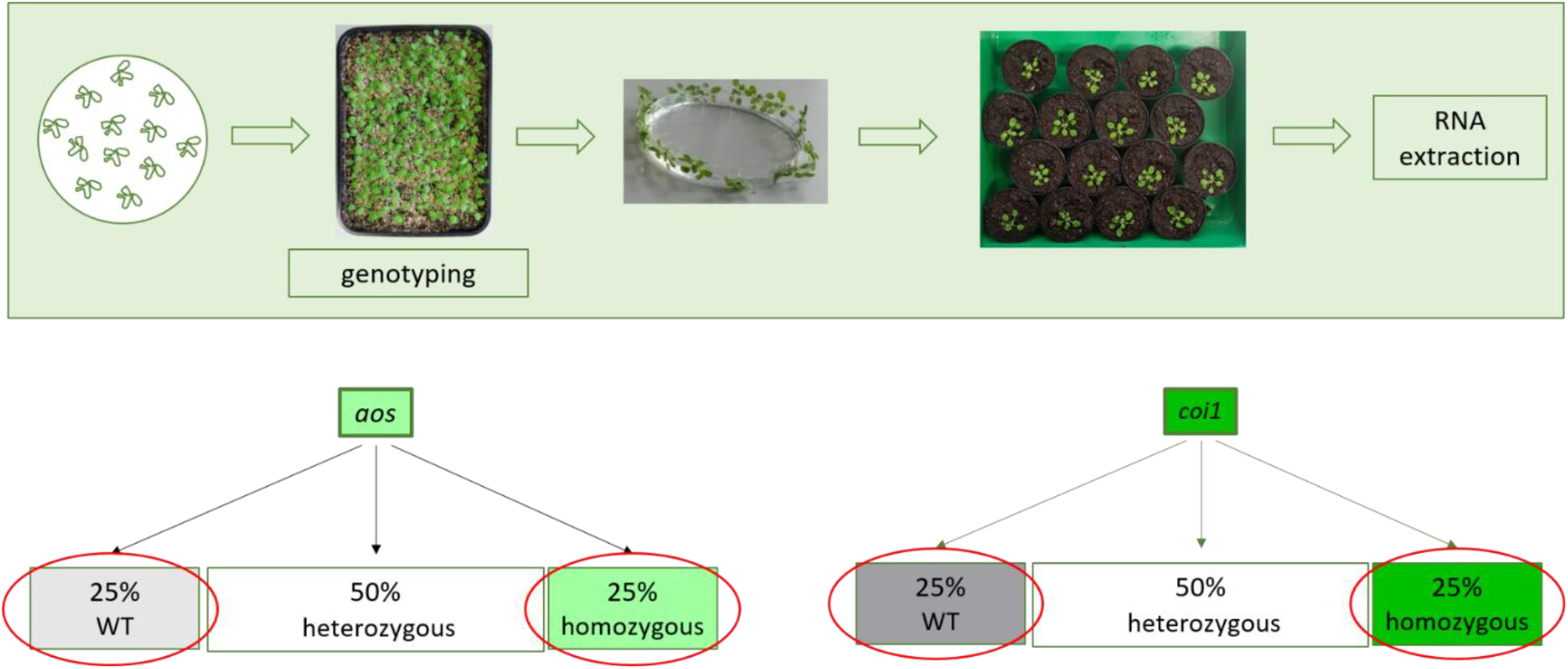
Experimental outline of plant treatments for RNAseq analysis. Plants were first grown on agar plates containing Murashige-Skoog-medium (MS) supplemented with 2% sucrose. After 14 days, they were carefully transferred onto a 1:1 mix of sand and twice-steamed soil on a thin layer of Seramis. Cultivation was continued for another 14 days under short-day conditions at a photon flux density of 120-140 μmol m^-2^ s^-1^. For genotyping, a single leaf was clipped from each plant during the first week of growth on the sand-soil mixture. For either mock treatment or fungal infection, plants were carefully up-rooted and roots were washed in tap water. Roots were then dipped in tap water as mock treatment or in a *V. longisporum* spore suspension for 45 minutes. Afterwards, plants were planted into individual pots containing twice-steamed soil and kept for a final 10 days in short day conditions at 120-140 μmol photons m^-2^ s^-1^. Root and shoot material of one single plant were harvested for one biological replicate.

**Fig. S2.**
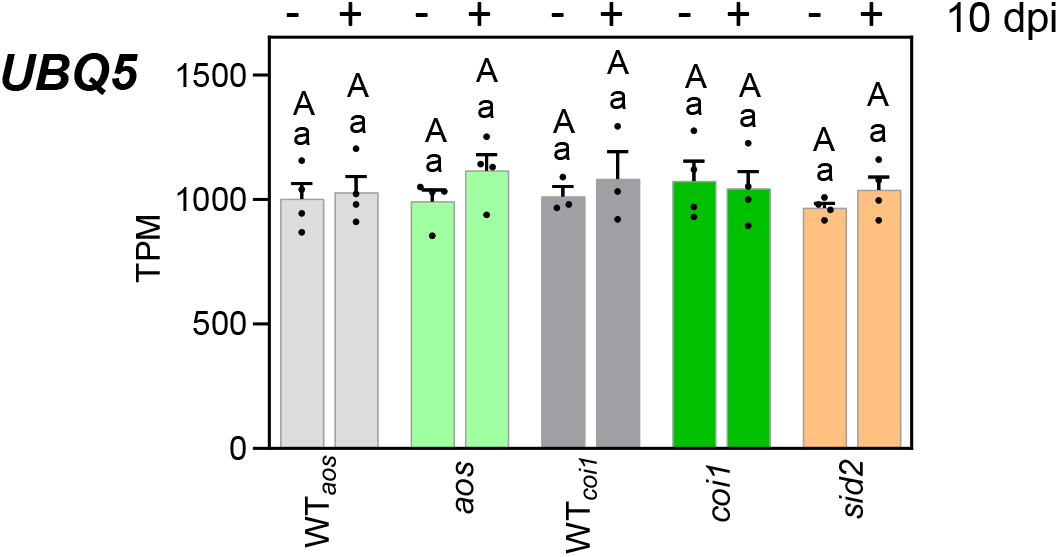
*UBQ5* is expressed independently of genotype and treatment. Relative *UBQ5* transcript levels as quantified by RNAseq analysis 10 days after mock treatment or inoculation with 1×10^6^ spores/mL eGFP-expressing *V. longisporum*. Bars are means of Transcripts Per Million (TPM) ± SEM of three to four biological replicates of each genotype, with each replicate representing twelve roots from one independent experiment. For statistical analysis logarithmic values were subjected to a two-way ANOVA analysis followed by Tukey’s multiple comparison test; lowercase letters denote significant differences within each genotype between mock and 10 dpi (*p* < 0.05), uppercase letters denote significant differences between genotypes subjected to the same treatment (*p* < 0.05). WT*_aos_* and WT*_coi1_* are the two wild-type lines obtained from the segregating offspring of heterozygous *aos* and *coi1* seeds.

**Fig. S3.**
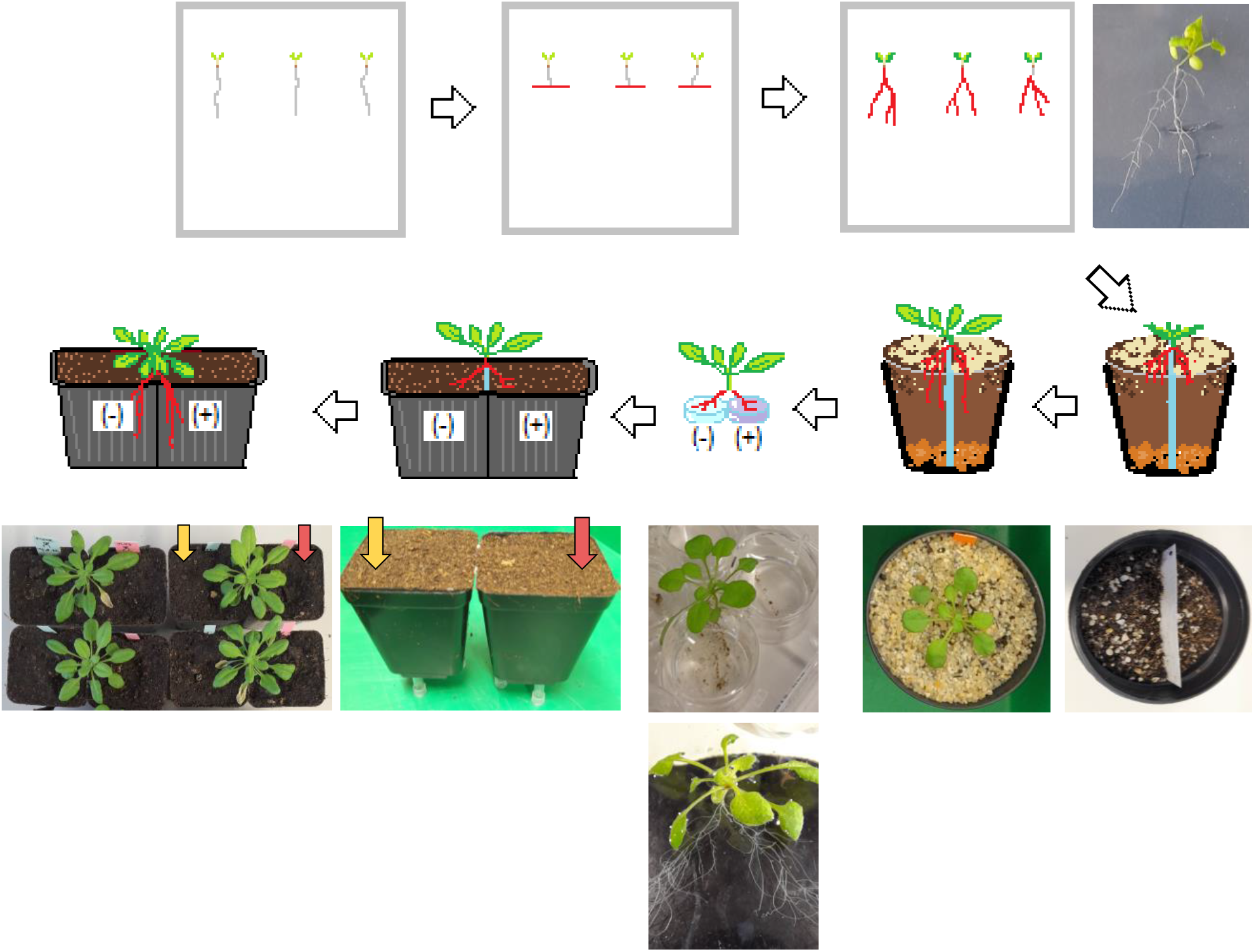
Experimental outline of the split-root experiment. Seedlings were grown vertically on Murashige-Skoog-medium (MS) supplemented with 2% sucrose. After 7 days, primary roots were cut below the first two lateral roots, and roots were allowed to grow for additional 14 days. Subsequently, each plant with two main roots of similar size was transferred onto a pot, in which two compartments were established with a small barrier made of cut overhead transparencies. After 14 days of cultivation on the 1:1 mix of sand and twice-steamed soil on a thin layer of Seramis (short-day conditions at a photon flux density of 120-140 μmol m^-2^ s^-1^), the two root systems were carefully placed into a 12 well microtiter plate containing either the spore suspension or tap water. Subsequently, plants were transferred to soil, with both root systems growing separately in two adjacent pots. After 12 days, roots of each pot were harvested separately.

**Fig S4.**
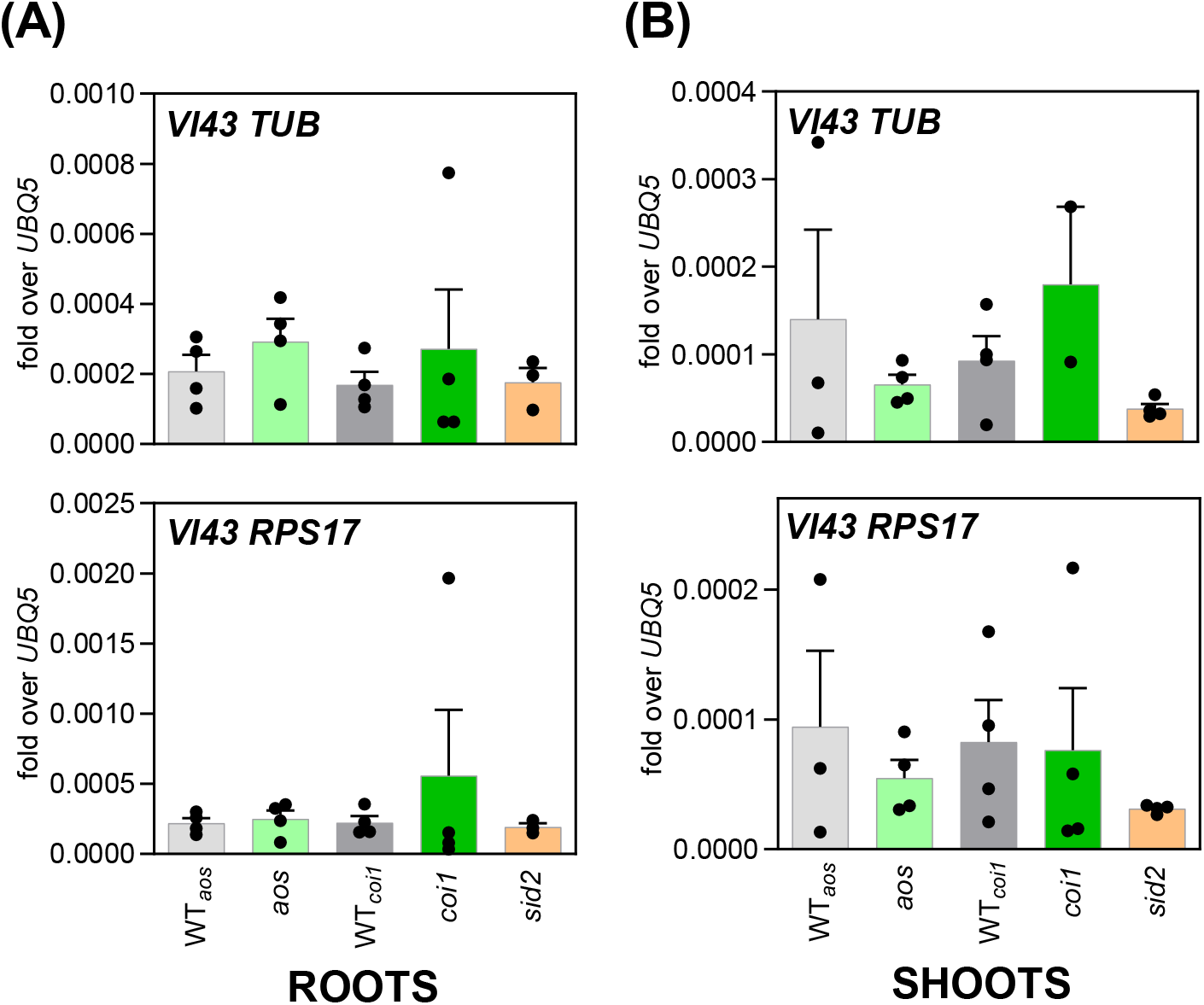
Detection of fungal transcripts in different plant genotypes after infection with *V. longisporum*. Fungal *TUBULIN (TUB)* and *RIBOSOMAL PROTEIN S17 (RPS17)* transcripts were quantified via qRT-PCR and normalized to the abundance of Arabidopsis *UBQ5* transcripts. RNA was extracted from **(A)** roots (same material as used for RNAseq analysis) or **(B)** shoots of WT*_aos_*, WT*_coi1_*, *aos, coi1* and *sid2* plants infected with *Verticillium longisporum* (10 dpi) from the same experiment. WT*_aos_* and WT*_coi1_* are the wild-types obtained from the segregating offspring of heterozygous *aos* and *coi1* seeds. Bars are means ± SEM of three to four biological replicates of each genotype, with each replicate representing twelve roots or shoots from one independent experiment. For statistical analysis, a one-way ANOVA was performed followed by Tukey’s multiple comparison test; no significant differences were observed (*p* < 0.05).

**Fig. S5.**
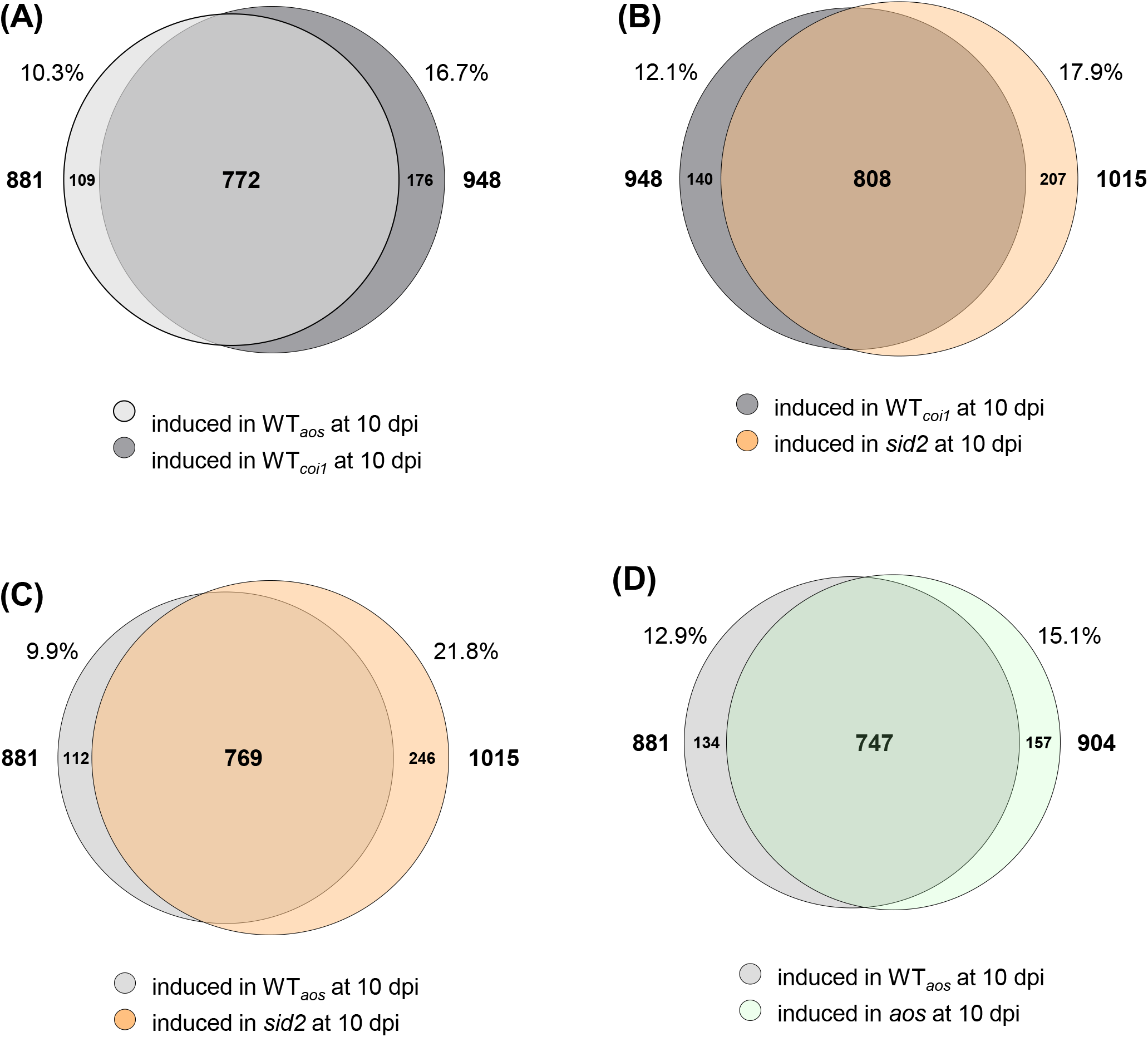
Venn diagrams of genes induced in WT*_aos_*, WT*_coi1_*, *aos* and *sid2* after infection with *V. longisporum.* Venn diagrams showing. **(A)** the overlap between genes induced in WT*_aos_* and WT*_coi1_* at 10 dpi (> 2-fold, *p* < 0.05), **(B)** the overlap between genes induced in WT*_aos_* and *aos* at 10 dpi (> 2-fold, *p* < 0.05), **(C)** the overlap between genes induced in WT*_aos_* and *sid2* at 10 dpi (> 2-fold, *p* < 0.05), **(D)** the overlap between genes induced in WT*_coi1_* and *sid2* at 10 dpi (> 2-fold, *p* < 0.05). Expression data was obtained by RNAseq analysis from root material 10 days after mock treatment or inoculation with 1×10^6^ spores/mL eGFP-expressing *V. longisporum*. Circles are drawn to scale with respect to the number of genes represented in each group. WT*_aos_* and WT*_coi1_* are the wild-types obtained from the segregating offspring of heterozygous *aos* and *coi1* seeds.

**Fig. S6.**
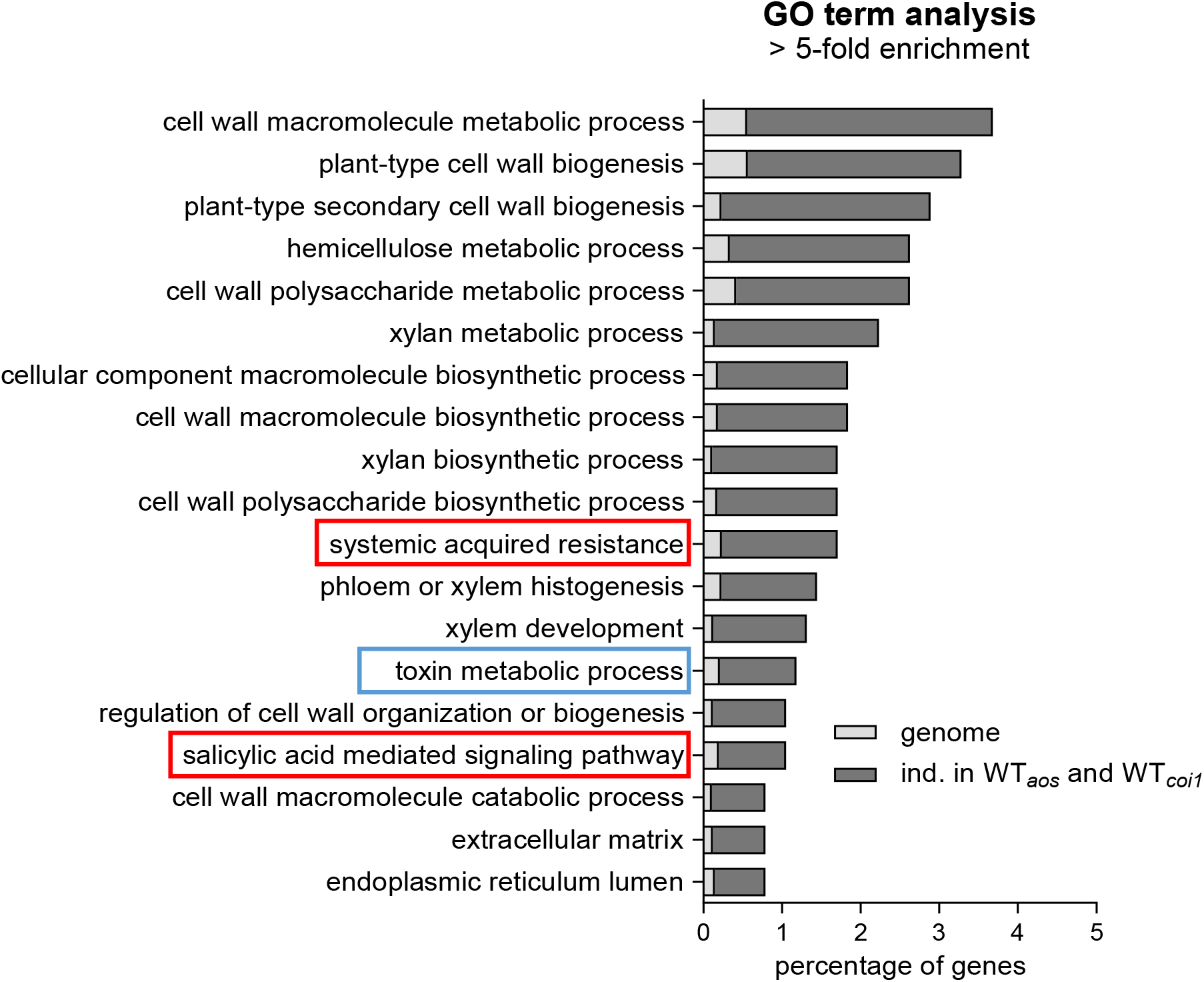
Gene Ontology (GO) term enrichment analysis of 772 genes significantly induced in roots of WT*_aos_* and WT*_coi1_* after infection with *V. longisporum*. Bars represent the percentage of genes found per GO term in the group of 772 genes induced at 10 dpi in WT*_aos_* and WT*_coi1_* (dark grey) and the percentage of genes representing the respective GO term found within the Arabidopsis genome (light grey). GO terms containing > 0.1% of genes of the genome and with > 5-fold enrichment against the genome are shown. Statistical analysis was performed using Fisher’s Exact test and False Discovery Rate (FDR) < 0.05. Boxes mark GO terms not related to xylem vessel formation.

**Fig. S7.**
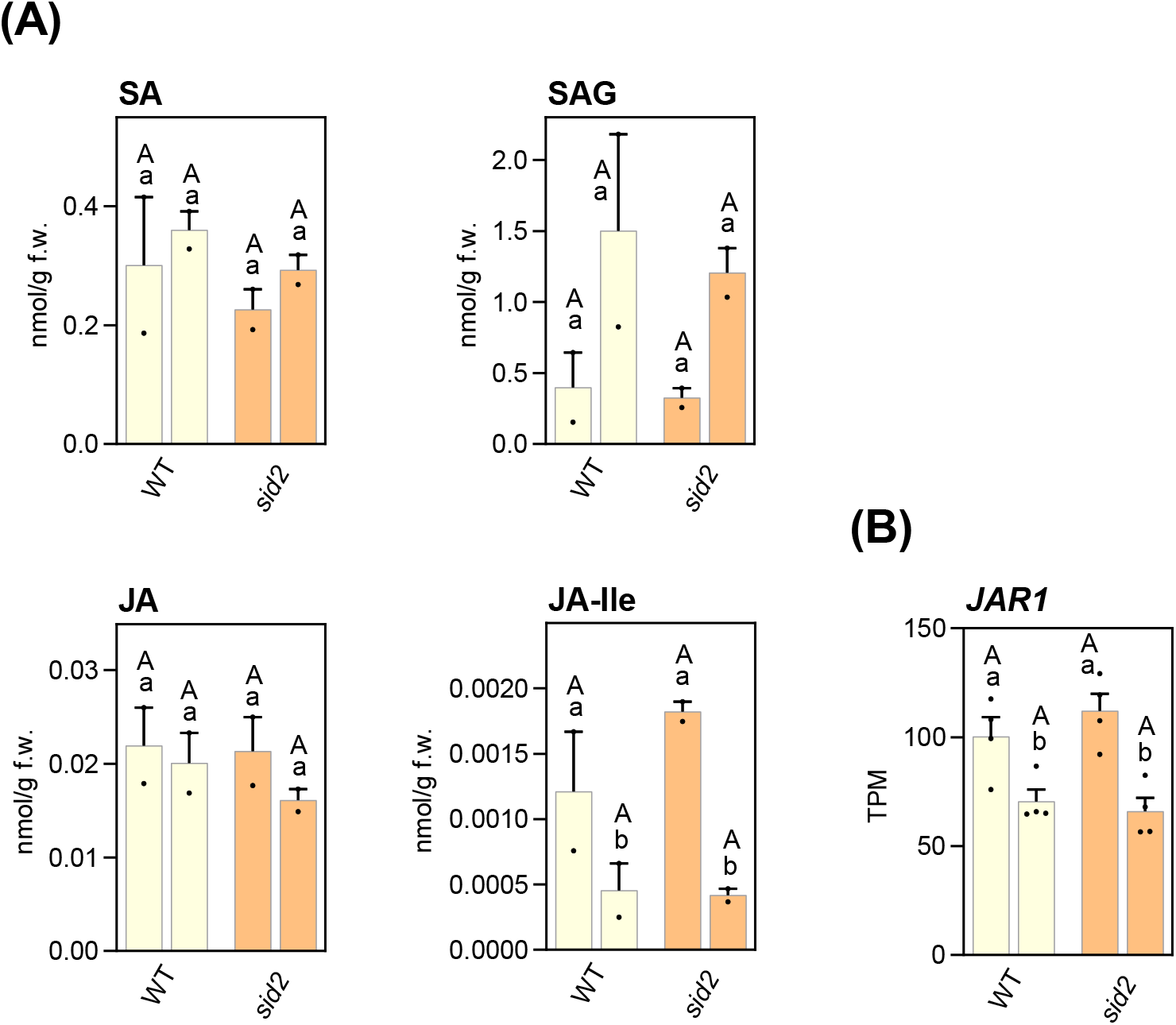
Defense hormones in *V. longisporum*-infected roots. **(A)** Phytohormone levels in roots at 10 days after mock treatment or infection with 1×10^6^ spores/mL *V. longisporum*. Eight to ten roots were pooled per sample. Bars are means ± SEM of two samples per genotype. **(B)** Relative *JAR1* transcript levels as quantified by RNAseq analysis 10 days after mock treatment or inoculation with 1×10^6^ spores/mL eGFP-expressing *V. longisporum*. Bars are means of Transcripts Per Million (TPM) ± SEM of four biological replicates of each genotype, with each replicate representing twelve roots from one independent experiment. Analysis was done as described in: YU, D., JANZ, D., ZIENKIEWICZ, K., HERRFURTH, C., FEUSSNER, I., CHEN, S. & POLLE, A. 2021. Wood Formation under Severe Drought Invokes Adjustment of the Hormonal and Transcriptional Landscape in Poplar. Int J Mol Sci, 22. For statistical analysis, a two-way ANOVA was performed followed by Tukey’s multiple comparison test; lowercase letters denote significant differences within each genotype between mock and 10 dpi (*p* < 0.05), uppercase letters denote significant differences between genotypes subjected to the same treatment (*p* < 0.05). For **(B)**, logarithmic values were used for the statistical analysis.

**Fig. S8.**
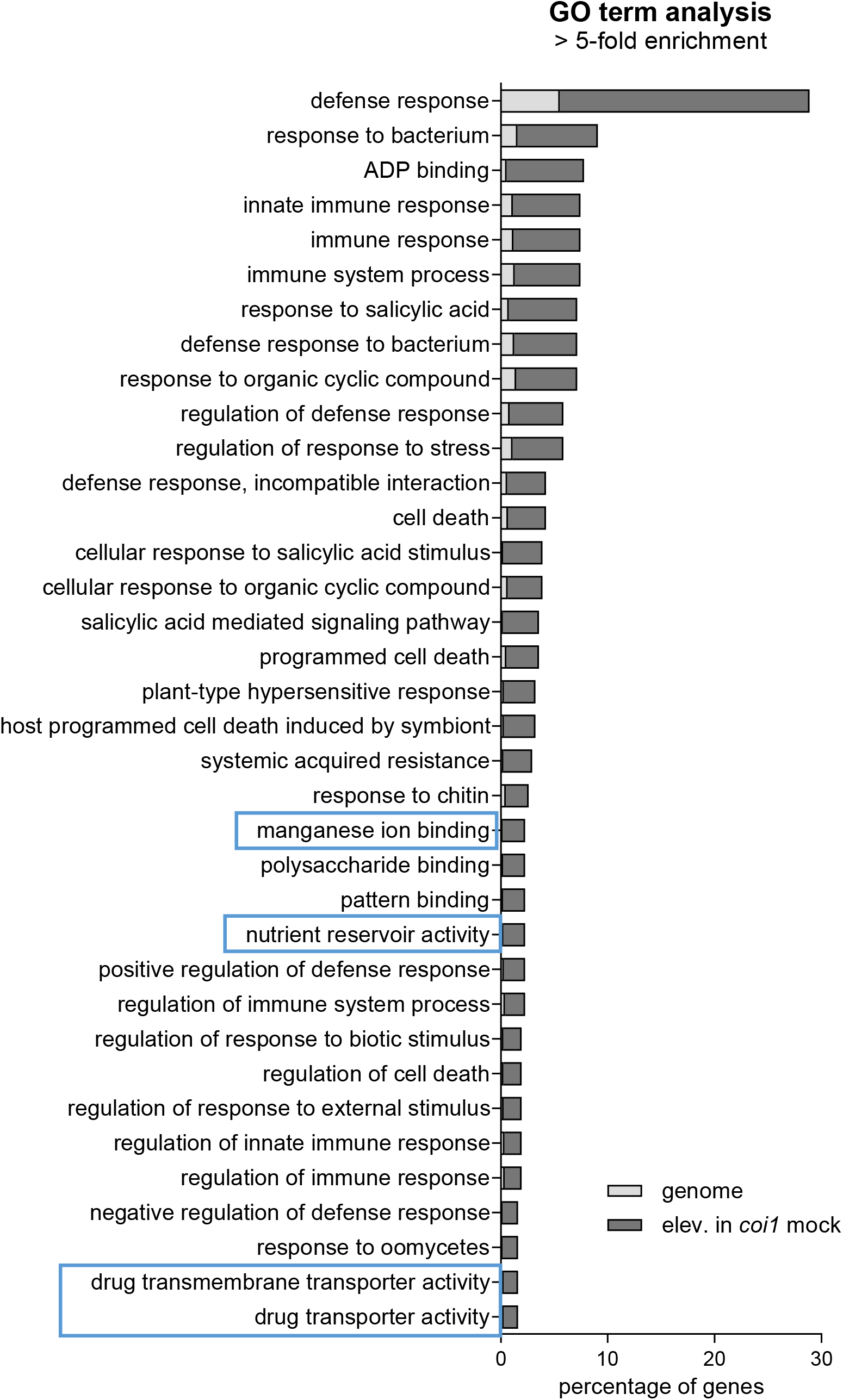
Gene Ontology (GO) term enrichment analysis of 316 genes significantly higher expressed in mock-treated *coi1* than in all other genotypes. Dark grey bars represent the percentage of genes found per GO term in the group of 316 genes significantly higher expressed at 10 days after mock treatment in *coi1* (dark grey) as in WT*_aos_*, WT*_coi1_*, *aos* and *sid2*; light grey bars represent the percentage of genes of the respective GO term found within the Arabidopsis genome. GO terms containing > 0.1% of genes of the genome and with > 5-fold enrichment against the genome are shown. Statistical analysis was performed using Fisher’s Exact test and False Discovery Rate (FDR) < 0.05. Boxes mark GO terms not related to defense responses.

**Fig. S9.**
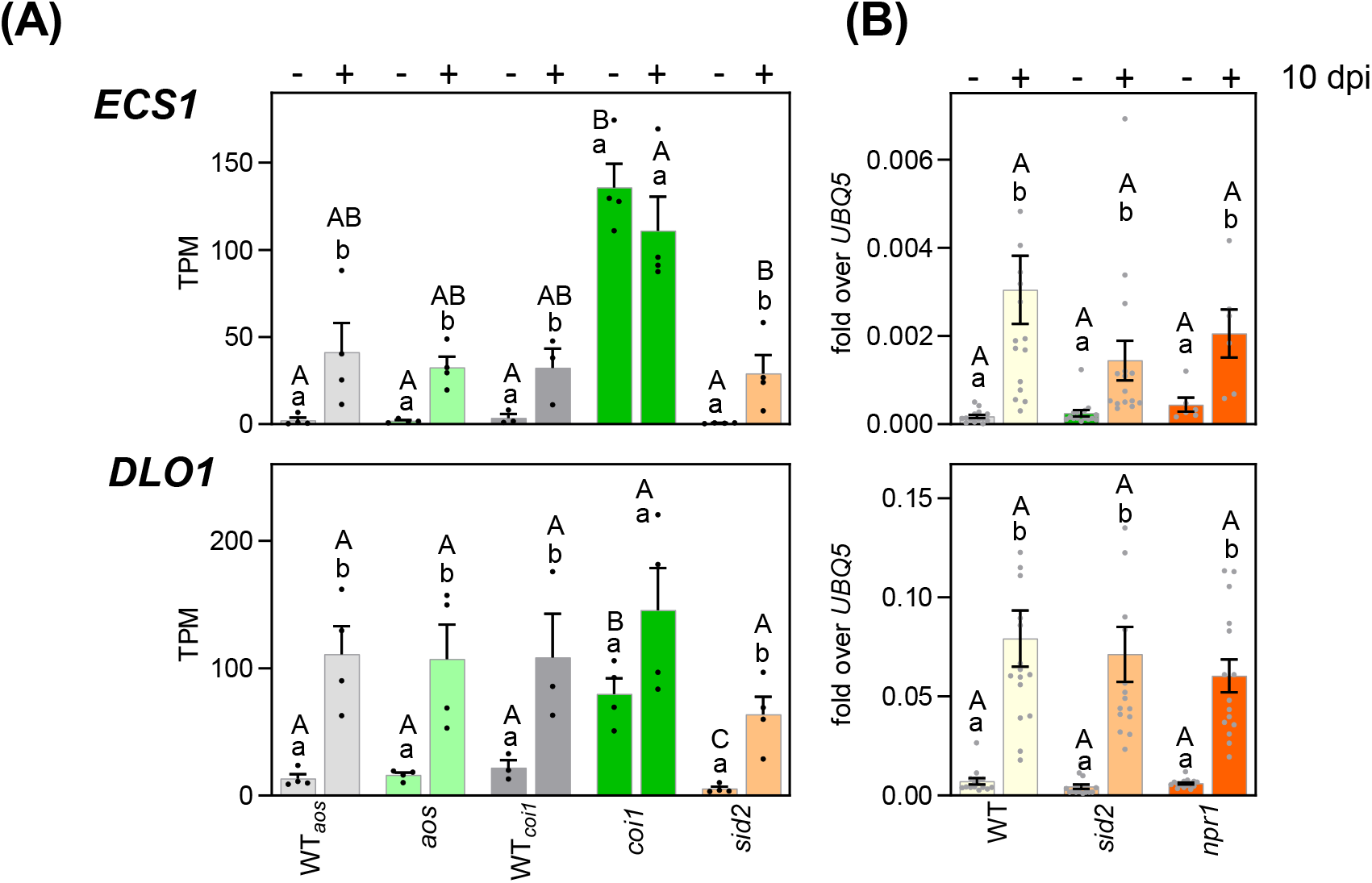
Expression analyses of *ECS1* and *DLO1* in different genotyes after *V. longisporum* infection. **(A)** Relative *ECS1* and *DLO1* transcript levels as quantified by RNAseq analysis 10 days after mock treatment or inoculation with 1×10^6^ spores/mL eGFP-expressing *V. longisporum*. Bars are means of Transcripts Per Million (TPM) ± SEM of three to four biological replicates of each genotype, with each replicate representing twelve roots from one independent experiment. **(B)** Relative *ECS1* and *DLO1* transcript levels in WT, *sid2* and *npr1*, measured by qRT-PCR. RNA was extracted from roots 10 days after mock treatment or infection with 1×10^6^ spores/mL *V. longisporum*. Bars are means ± SEM of thirteen to sixteen roots per genotype. For statistical analysis logarithmic values were subjected to a two-way ANOVA analysis followed by Tukey’s multiple comparison test; lowercase letters denote significant differences within each genotype between mock and 10 dpi (*p* < 0.05), uppercase letters denote significant differences between genotypes subjected to the same treatment (*p* < 0.05). WT*_aos_* and WT*_coi1_* are the two wild-type lines obtained from the segregating offspring of heterozygous *aos* and *coi1* seeds.

**Fig. S10.**
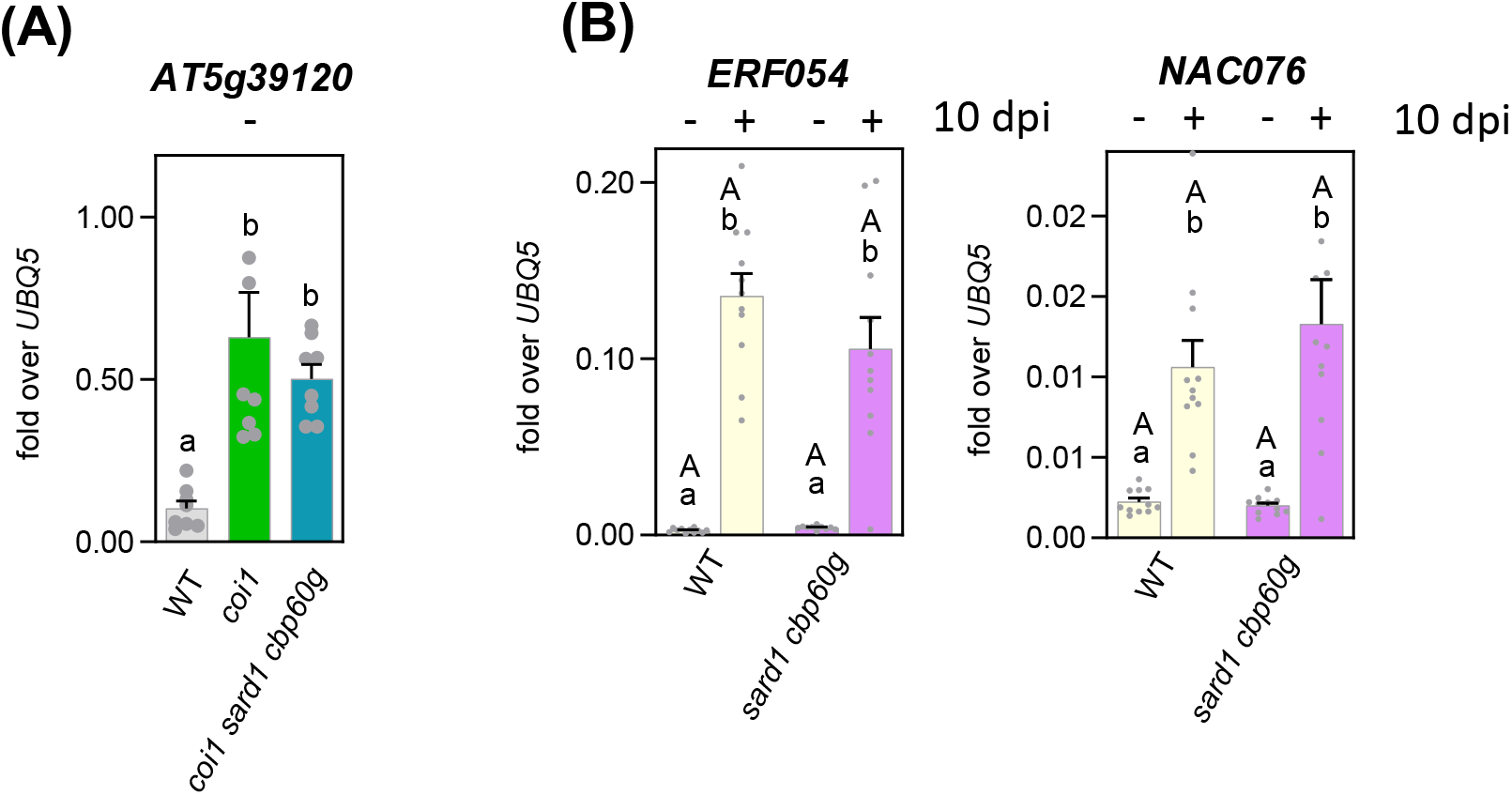
Selected genes of the ‘*coi1^up^*’ and ‘*Vl^ind^*’ clusters are not regulated by SARD1/CBP60g. **(A)** *AT5G39120* (a RmlC-like cupins superfamily protein, belonging to the ‘*coi1^up^*’ cluster) transcript levels, measured by qRT-PCR. RNA was extracted from roots 10 days after mock treatment. Bars are means ± SEM of six to eight roots per genotype. **(B)** *ERF054* and *ANAC076* (which belong to the ‘*Vl^ind^*’ cluster) transcript levels, measured by qRT-PCR. RNA was extracted from roots 10 days after mock treatment or infection with 1×10^6^ spores/mL *V. longisporum*. Bars are means ± SEM of ten to eleven roots per genotype. For statistical analysis with logarithmic values, either a one-way (A) or two-way ANOVA was performed followed by a Tukey’s multiple comparison test; lowercase letters denote significant differences within each genotype between mock and 10dpi (*p* < 0.05), uppercase letters denote significant differences between genotypes subjected to the same treatment (*p* < 0.05).

**Fig. S11.**
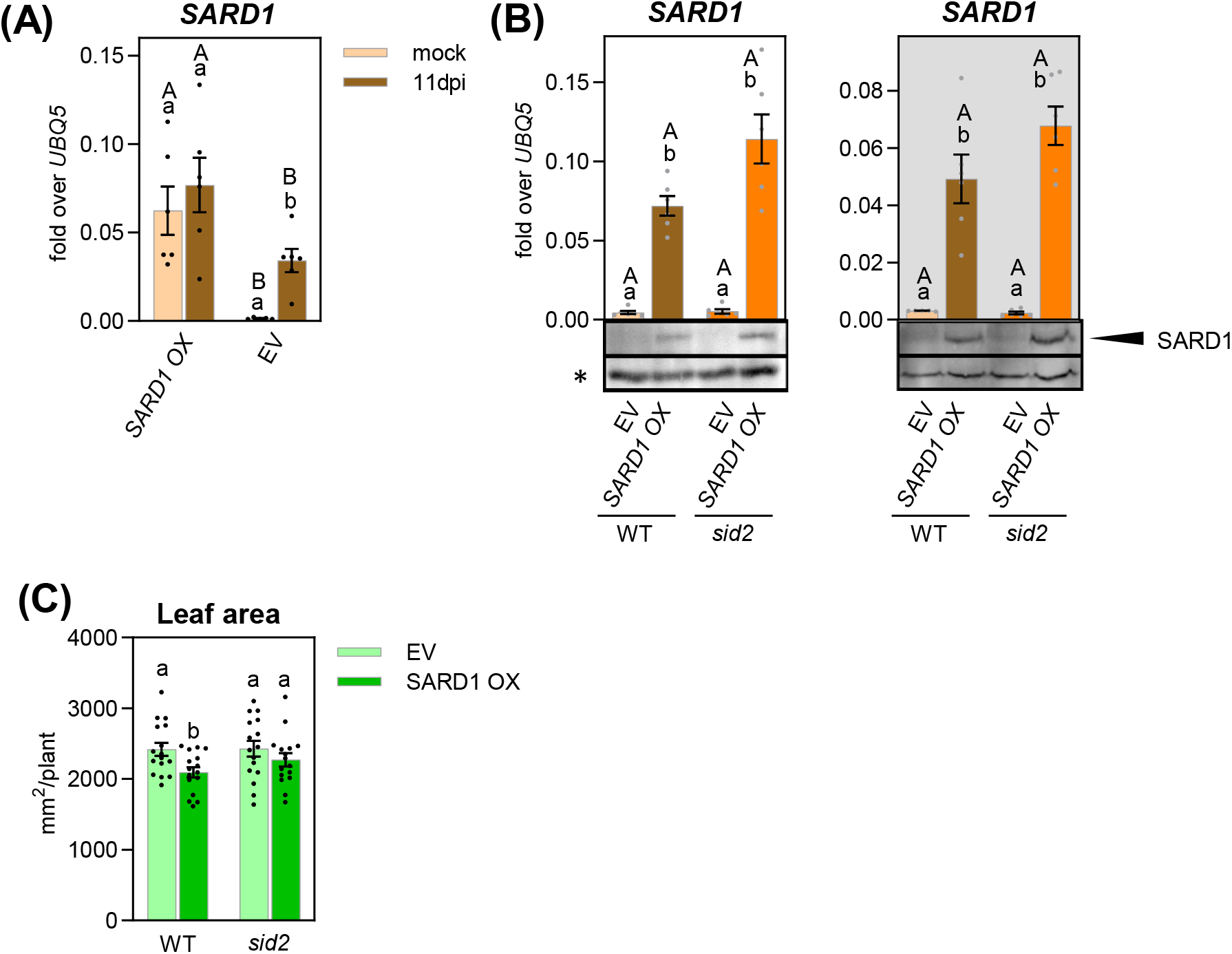
Characterization of *SARD1* overexpressing lines. **(A)** Relative SARD1 transcript levels, measured by qRT-PCR. RNA was extracted from roots 10 days after mock treatment or infection with 1×10^6^ spores/mL *V. longisporum*. Bars are means ± SEM of ten to eleven roots per genotype. **(B)** Relative *SARD1* transcript levels, measured by qRT-PCR. RNA was extracted from shoots or roots (grey background of plotting area) at 10 days after mock treatment of *SARD1* overexpression lines (*SARD1 OX*) and empty vector (EV) controls in WT and *sid2*. Bars are means ± SEM of three to six roots or shoots per line. Inserts: Western blot of protein extracts obtained from roots and shoots of *SARD1* overexpression lines (*SARD1 OX*) and empty vector (EV) controls in WT and *sid2*. Per lane, six roots or three shoots were pooled from each line. C-terminally 3xHA-StrepII tagged SARD1 protein levels were detected using an anti-HA antibody. * depicts an unspecific band shown as loading control. **(C)** Leaf area of *SARD1* overexpression lines (*SARD1 OX*) and empty vector (EV) controls in WT and *sid2*. Values are from 16 plants per genotype. For statistical analysis with logarithmic (transcript) or linear (leaf area) values, a two-way ANOVA followed by Tukey’s multiple comparison test was performed; lowercase letters denote significant differences within each genotype (WT and *sid2*) between EV and SARD1 OX **(a, c)** or mock and 10 dpi (*p* < 0.05) **(B),** uppercase letters denote significant differences between genotypes transformed with the same construct (EV and SARD1 OX) **(a, c)** or the same treatment **(B)** (*p* < 0.05). Bars represent means ± SEM

**Fig. S12.**
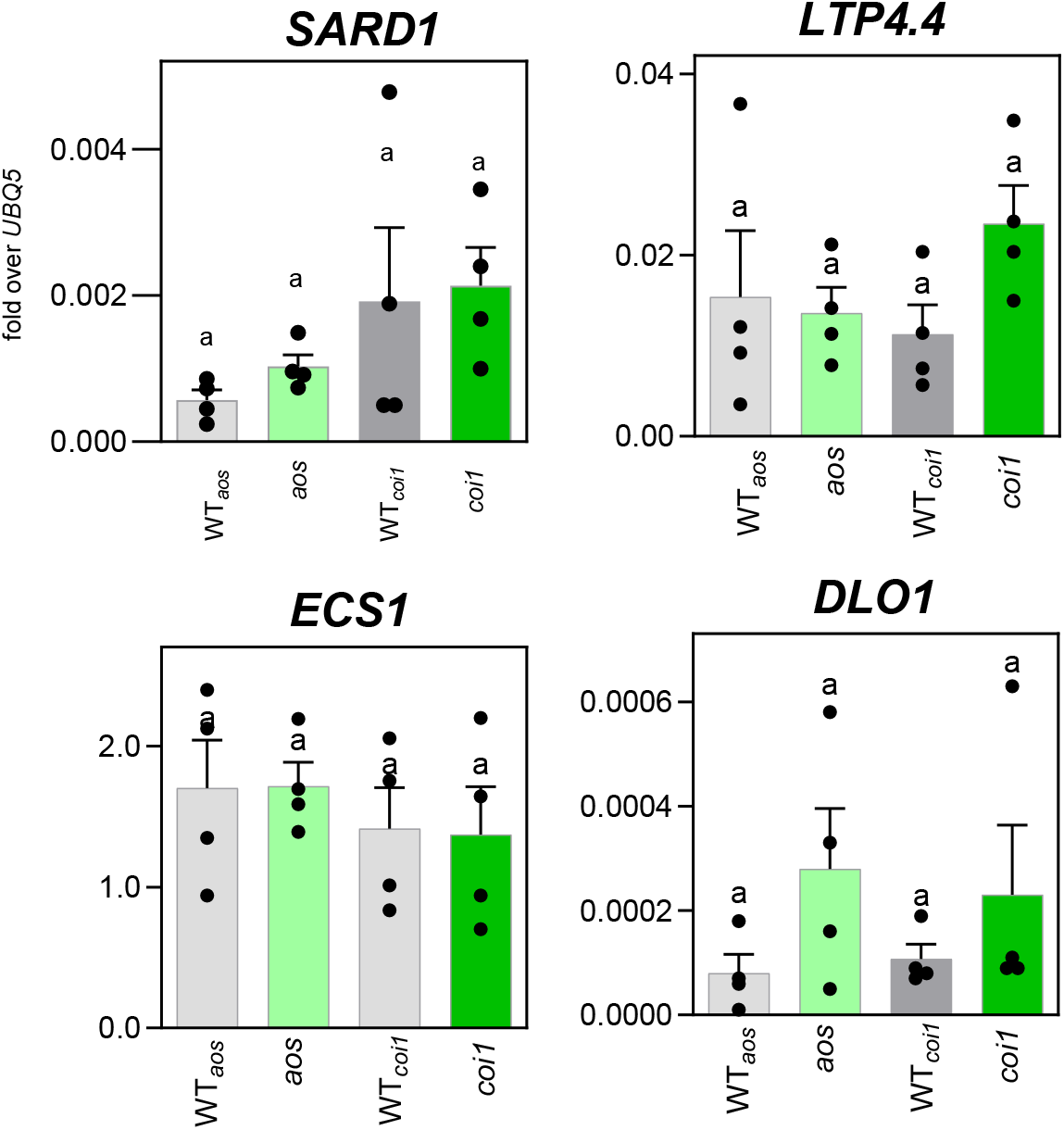
COI1 is not a constitutive repressor of target genes in shoots. *SARD1, LTP4.4, ECS1* and *DLO1* transcript levels, measured by qRT-PCR. RNA was extracted from whole rosettes 10 days after mock treatment from the same plants the roots of which were subjected to the RNAseq analysis. Bars are means ± SEM of four replicates, each from twelve rosettes per genotype. For statistical analysis, a one-way ANOVA was performed followed by Tukey’s multiple comparison test; no significant differences were observed (*p* < 0.05).

**Fig. S13.**
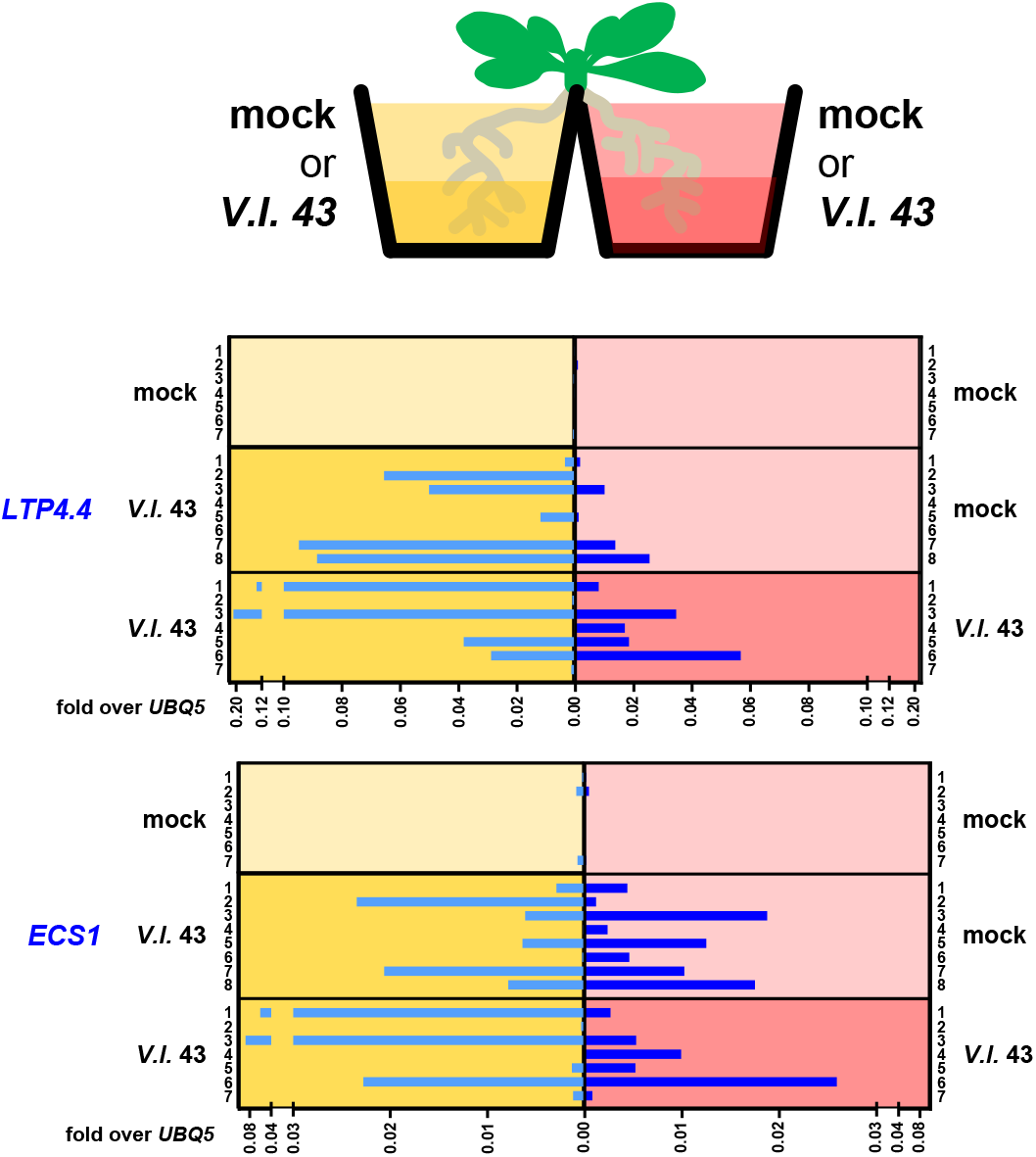
Expression analysis of *V. longisporum*-induced genes in physically separated roots. The two physically separated root systems were independently treated (mock/mock; *V. longisporum*/mock and *V. longisporum*/*V. longisporum*) RNA was extracted at 12 dpi. Relative expression of selected marker genes was determined by real time PCR, and values of the two root systems of each individual plant were plotted next to each other. The same RNA as shown in Figure 6 was used for the analysis

**Fig. S14.**
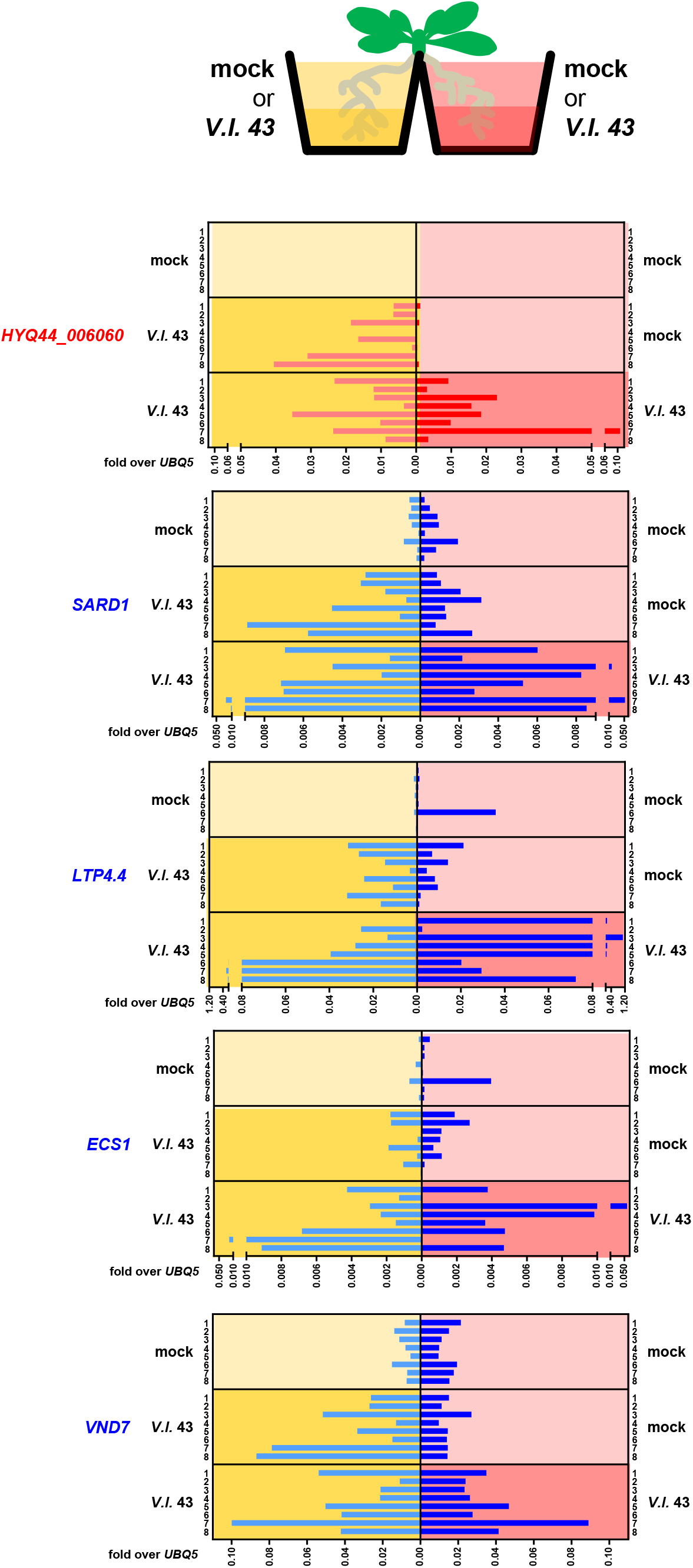
Expression analysis of *V. longisporum*-induced genes in physically separated roots (Experiment #2). The two physically separated root systems were independently treated (mock/mock; *V. longisporum*/mock and *V. longisporum*/*V. longisporum*) RNA was extracted at 12 dpi. Relative expression of selected marker genes was determined by real time PCR, and values of the two root systems of each individual plant were plotted next to each other.

## Author contributions

L.U. performed most of the experiments and analysed most of the data; J.S. performed experiments for RNAseq; C.T. and analysed the RNAseq data; C.T. and C.G. designed and supervised the research; L.U. and C.G. wrote the manuscript with input from all authors.

## Acknowledgements

We thank Anna Hermann, Katharina Dworak and Ronald Scholz for excellent technical assistance and Natalie Leutert for help with characterisation of transgenic *SARD1 OX* lines. We also thank the Next Generation Sequencing-Integrative Genomics Core Unit (NIG), Institute of Human Genetics, at the University Medical Centre Göttingen (UMG), Germany, for performing the RNAseq analysis and Goettingen Center for Molecular Biosciences (GZMB), Service Unit for Metabolomics and Lipidomics, Goettingen, Germany, for phytohormone quantifications.

## Data availability statement

The data that support the findings of this study are available from the corresponding author upon reasonable request.

